# Fast Medical Image Auto-segmentation for Bleeding Gastric Tissue Detection based on Deep DuS-KFCM Clustering

**DOI:** 10.1101/2021.12.22.473941

**Authors:** Xian-Xian Liu, Gloria Li, Wei Luo, Juntao Gao, Simon Fong

## Abstract

**Background:** Detection and classification of gastric bleeding tissues are one of the challenging tasks in endoscopy image analysis. Lesion detection plays an important role in gastric cancer (GC) diagnosis and follow-up. Manual segmentation of endoscopy images is a very time-consuming task and subject to intra- and interrater variability. Accurate GB segmentation in abdominal sequences is an essential and crucial task for surgical planning and navigation in gastric lesion ablation. However, GB segmentation in endoscope is a substantially challenging work because the intensity values of gastric blood are similar to those of adjacent structures.

**Objective:** In this paper the idea is to combine two parts: Neural Network and Fuzzy Logic--Hybrid Neuro-Fuzzy system. The objective of this manuscript is to provide an efficient way to segment the gastric bleeding lesion area. This work focuses on design and development of an automated diagnostic system using gastric bleeding cancer endoscopy images.

**Methods:** In this paper, a coarse-to-fine method was applied to segment gastric bleeding lesion from endoscopy images, which consists of two stages including rough segmentation and refined segmentation. The rough segmentation is based on a kernel fuzzy C-means algorithm with spatial information (SKFCM) algorithm combined with spatial gray level co-occurrence matrix (GLCM) and the refined segmentation is implemented with deeplabv3+ (backbone with resnet50) algorithm to improve the overall accuracy.

**Results:** Experimental results for gastric bleeding segmentation show that the method provides an accuracy of 87.9476% with specificity of 96.3343% and performs better than other related methods.

**Conclusions:** The performance of the method was evaluated using two benchmark datasets: The GB Segmentation and the healthy datasets. Then use the gastric red spots (GRS) dataset to do the final test to verify weak bleeding symptoms. Our method achieves high accuracy in gastric bleeding lesion segmentation. The work describes an innovative way of using GLCM based textural features to extract underlying information in gastric bleeding cancer imagery. Modified deep DuS-KFCM endoscopy image segmentation method based on GLCM feature, The experimental results shown to be effective in image segmentation and has good performance of resisting noise, segmentation effect more ideal.

## 1.1. Introduction

Lesion detection plays an important role in early gastric cancer (EGC) diagnosis and follow-up. Manual segmentation of endoscopy images is a very time-consuming task and subject to intra and inter-rater variability. Nowadays, the world is suffering dissimilar kind of gastric diseases. Generally, the stomach is the majority dangerous organ in the digestive system, The overall survival for early gastric cancer (GC) either managed endoscopically or surgically may exceed 96% [1], whereas the majority of cancer cases in the western world are diagnosed at late stages, and the cumulative 5-year survival is ranging 20–40% [2]. Separation of healthy people from affected patients is the prime objective; this is possible only if diagnosis is early.

Analyzing and processing of endoscopy gastric bleeding lesion areas are the most challenging and upcoming field. The image segmentation is entailed with the division or separation of the image into regions of similar features. Numerous researchers have worked and developed methods for solving cancer problems by using medical image segmentation. The task of image segmentation is to divide an image into a number of non-overlapping regions, which have same characteristics such as gray level, color, tone, texture, etc. A lot of clustering-based methods have been proposed for image segmentation. The segmentation & lesion detection approaches fall mainly under 3 categories. These are as follows: 1) Thresholding approaches, 2) Region growing approaches, 3) Histogram based approaches, 4) Clustering approaches. Several authors suggested various algorithms for segmentation.

Jaskirat Kaur, Sunil Agrawal & Renu Vig.’s paper [3] presented thresholding and edge detection being one of the important aspects of image segmentation comes prior to feature extraction and image recognition system for analyzing images. It helps in extracting the basic shape of an image, overlooking the minute unnecessary details. In this paper using image segmentation (thresholding and edge detection) techniques different geo-satellite images, medical images and architectural images are analyzed. To quantify the consistency of results, error measure is used.

Prastawa[4] developed tumor segmentation and statistical classification of brain MR images using an atlas prior. There are few challenges associated with atlas-based segmentation. Atlas-based segmentation requires manual labeling of template MRI. Such issues with atlas-based tumor segmentation can be mitigated by devising complementary techniques to aid tumor segmentation.

Image segmentation plays a crucial role in many areas of medical imaging. We have implemented a solution for segmenting image from the article. We developed a new method for automatic lesion segmentation using gastric bleeding (GB) and gastric red spots (GRS) images. The approach is based on three main steps, pre-processing of the data (which is not covered by the model). The method has been validated using real benign cases acquired with different scanners. We aimed to develop a computer aided model in order to automatically detect and segment the GB Lesions in endoscopy images, and to provide a quantitative description and a qualitative description for each patient. The proposed method was applied to segment the bleeding lesions for the whole dataset of abdominal gastric bleeding images.

However, these methods based on feature extraction have weaknesses. For example, an image contains other stomach tissue with similar blood color. When the area with abundant capillaries in it along with the lesion may cause misclustering. To overcome these weaknesses and to enhance the performance, it is difficult to determine the best centroids clusters among other clusters using SKFCM for segmenting object. since it provides the global information about the image in terms of distribution of various color components. The drawback of using SKFCM model is that the color information contained in the three channels is highly correlated. In order to address this problem, we proposed a dual spatial kernel fuzzy C-Means clustering algorithm based on color thresholding and space feature vectors,which is called a novel fuzzy set algorithm, which is used for the segmentation of the lesion region. Using spatial information from SKFCM’s color space and GLCM, the work proposed improves the clustering process. then the neuro-fuzzy system combines the learning power of Artificial Neural Network system and explicit knowledge representation of fuzzy inference system. A novel deep dual spatial kernelized fuzzy C-means algorithm with application in medical image segmentation facilitates bleeding area of gastric detection.

The image processing method presented in this manuscript, dual spatially kernelized constrained fuzzy c-means (DuS-KFCM) allows an image segmentation fuzzy region, inspired by the spatial kernelized constrained fuzzy c-means method (SKFCM), but using a GLCM texture analysis induced by compression from normal and abnormal (kernelized constrained deep fuzzy clustering based on dual spatial information), and a consideration of the neighborhood by the introduction of color space and GLCM feature vectors dual spatial constraints in the objective function of FCM.

The structure of the manuscript is as follows: in Section 2, identified work is described within the context of our work; in Section 3, using this sect. to specify the main contribution of the manuscript; In sect.4, The brief explanation about the proposed method is described in the section. our work is explained; Section 5 describes the proposed algorithm DuS-KFCM and the experimental result is given. in section 6, the proposed method deep DuS-KFCM is depicted, and experimental results are presented in Section 7; In sect.8, the findings and discussion appear; with the focus on three aspects: 1) a discussion related to evaluation indicators; and finally, we discuss the limitations of our work and conclude our work with some possible extension in the future in Sect. 9.

## 2. Identified Tasks

The novel proposed method consists of three phases:

In Phase 1, The endoscopy images obtained for gastric bleeding cancer diagnosis are preprocessed by first converting them into grayscale images, these raw images are then converted into binary images, after which the unwanted part is removed. Preprocessing in GB lesion particularly consists of delineation of lesion from the background and bleeding border extraction.

Segmented binary stack is basically the separation of a region of interest (ROI) from the background of the image. ROI is the part of the image that we want to use. In the case of cancerous images, we need the lesion part to extract the features from the diseased part, which is used in the pre-processing phase to improve image quality.

In Phase 2, After passing through the stages of preprocessing and image segmentation, there awaits post-processing where the task is to grab features. color space and statistical features are computed from each segmented image. The extracted characteristics are serially fused, and on addition of a CFS, redundant features are eliminated. Each vector of optimized functionality is forwarded to the GB clustering. the bleeding region is segmented using a DuS-KFCM method.

In Phase 3, Deep learning has the advantage of generating directly from raw images the high-level feature representation. In addition to DuS-KFCM, deep learning is also being used in parallel, for fuzzy stack feature extraction and image recognition. For example, DeepLabV3+ (backbone with Resnet50) neural networks have been able to detect lesion with promising performance. Deeplav3 serves as the bottleneck of the fuzzy clustering module used to segment the actual lesions of gastric lesion.

In Phase 4, To test these algorithms, there are publicly available datasets. These include GRS dataset for gastric bleeding cancer testing.

## 3. Main contributions of this paper

The most significant novel contributions with respect to the available state of the art of this article is summarized as follows:

The clustering segmentation method developed in this manuscript combines the spatial kernel fuzzy C-means technique with the spatial gray level co-occurrence matrix (GLCM) method and it also takes in account the dual spatial information (color space & statistical method of examining texture that considers the spatial relationship of pixels). This combination results in a novel dual spatial kernel fuzzy C-means clustering for gastric bleeding segmentation of digital images.

A robust segmentation methodology has been developed using mainly color clustering and texture feature, which is able to separate three regions of interest: GB, normal tissue and external region of gastric (non-gastric zone).

An innovative feature extraction method has been proposed in this manuscript. This method quantitatively calculates computes color-texture features. The novelty lies in that the color-texture information is obtained through the extraction of the gray level contrast, gray-tone linear dependencies, the complex degree of the texture and the evenness distribution degree of the texture features on different color components.

## 4. Proposed Methodology

### 4.1. Overview of the Proposed System

Gastric bleeding lesion segmentation seeks to separate healthy tissue from lesions regions. This is an essential step in diagnosis and treatment planning in order to maximize the likelihood of successful treatment. Due to the slow and tedious nature of manual segmentation, computer algorithms that do it faster and accurately are required. Because of the unpredictable appearance and shape of a bleeding area, segmenting gastric bleeding lesions from imaging data is one of the most challenging tasks in medical image analysis.

We have collected our data from the GitHub Data-Open-Access4PLoS-One datasets [1]. This folder contains brain images of 54 Patients. A novel algorithm is proposed in this paper to facilitate medical image segmentation. It is able to directly evolve from the initial segmentation by deep dual spatial fuzzy clustering.

Figure.1 illustrates the block diagram of the main stages of proposed model. The proposed architecture of fuzzy GB detection shows these steps.

**Figure 1.**
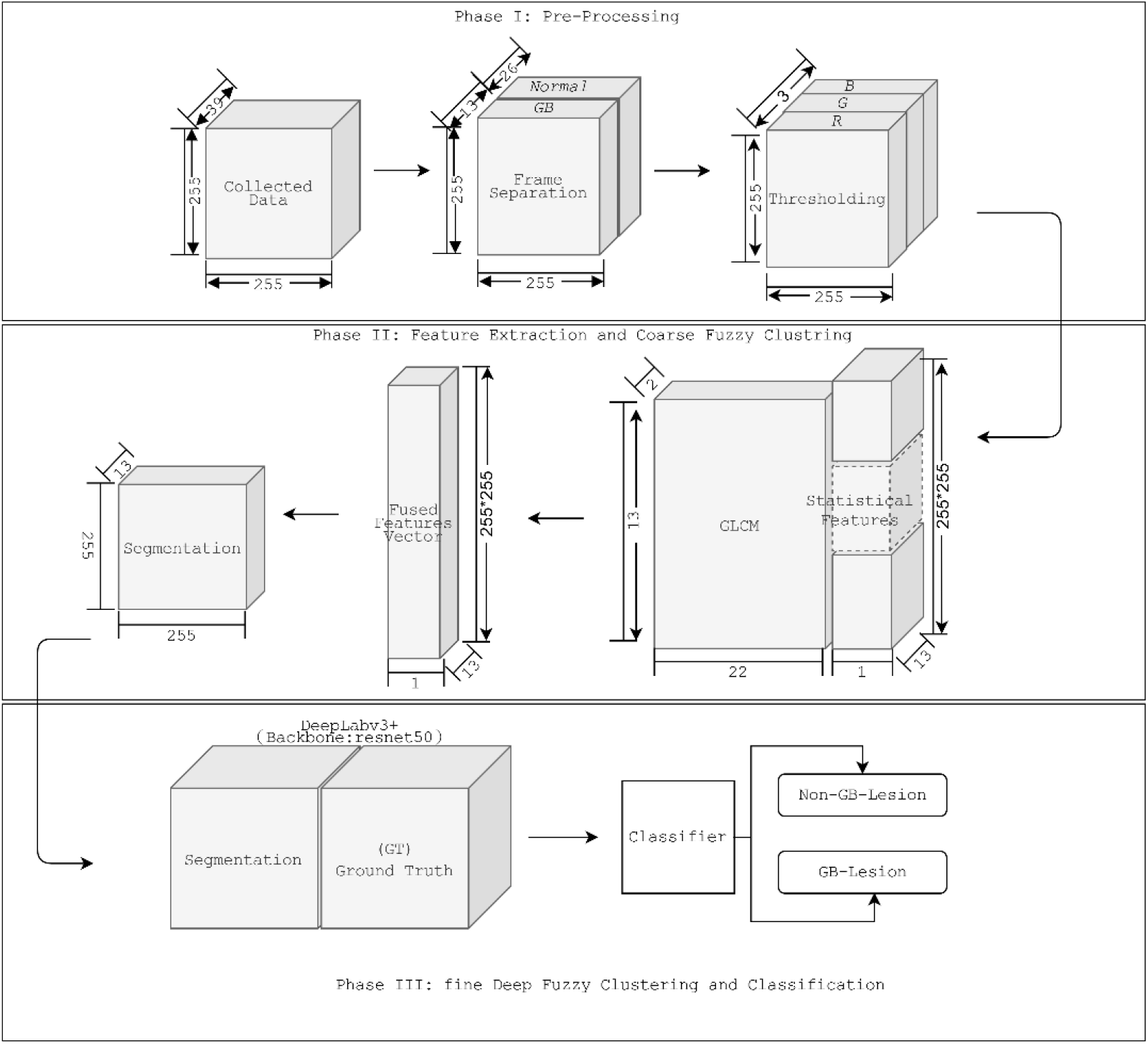
Graphical abstract: The block diagram of the proposed GB segmentation method.

Publicly available Since the ground truths for the challenges only included the segmentation mask, to produce the ground truth for ROI, we circumscribed a rectangle bounding box on the segmentation mask as shown in the Figure 2.

**Figure 2.**
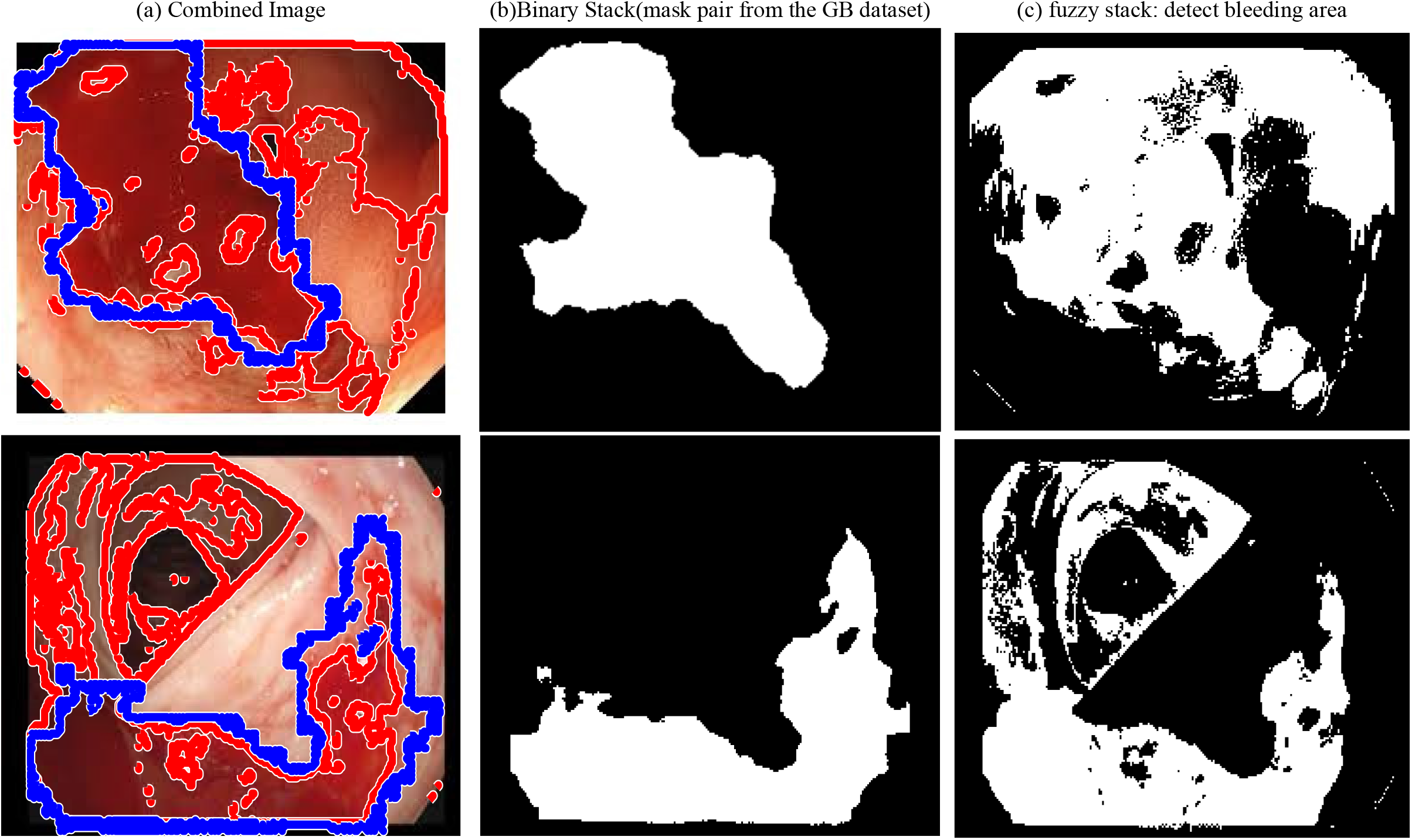
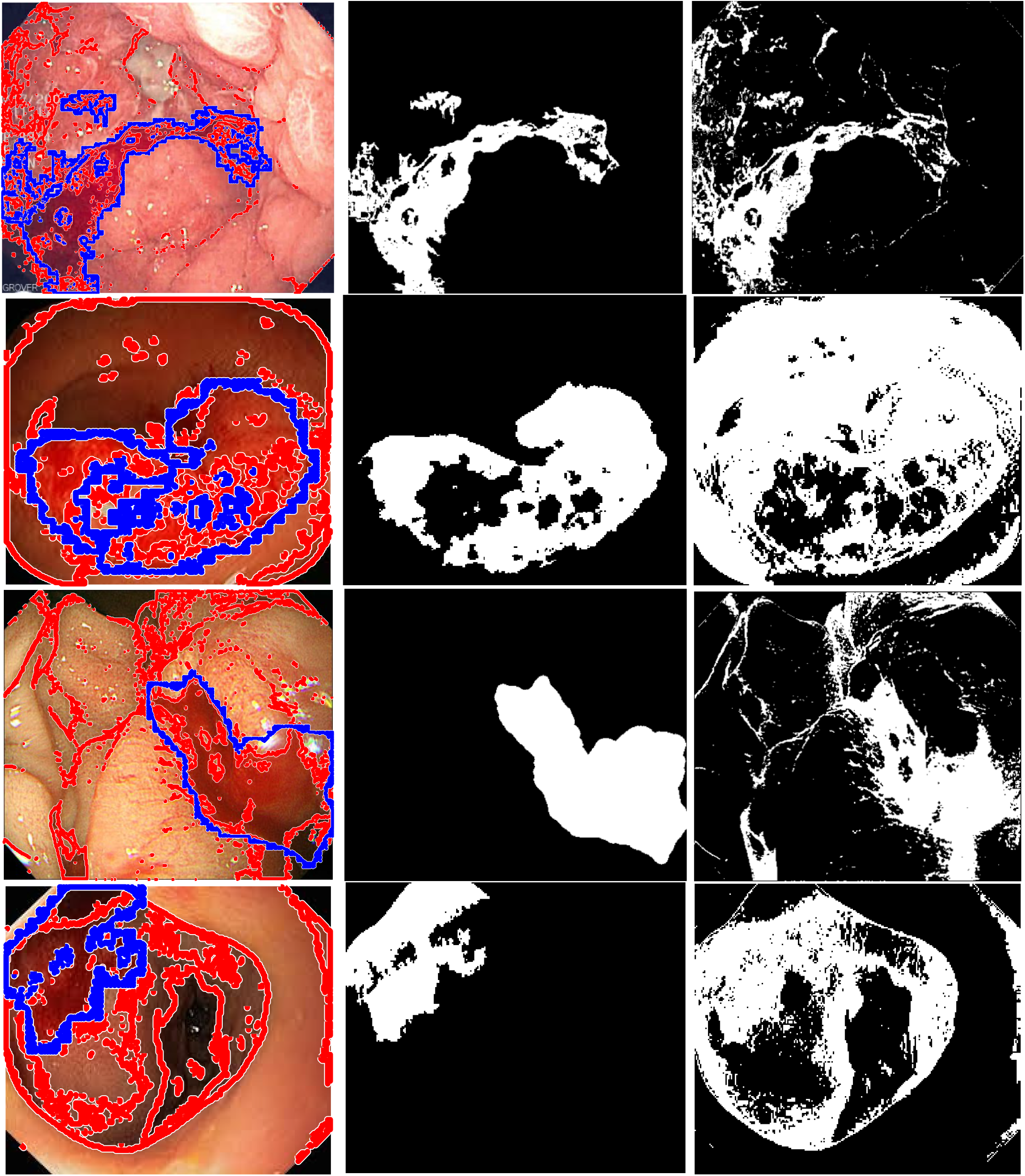
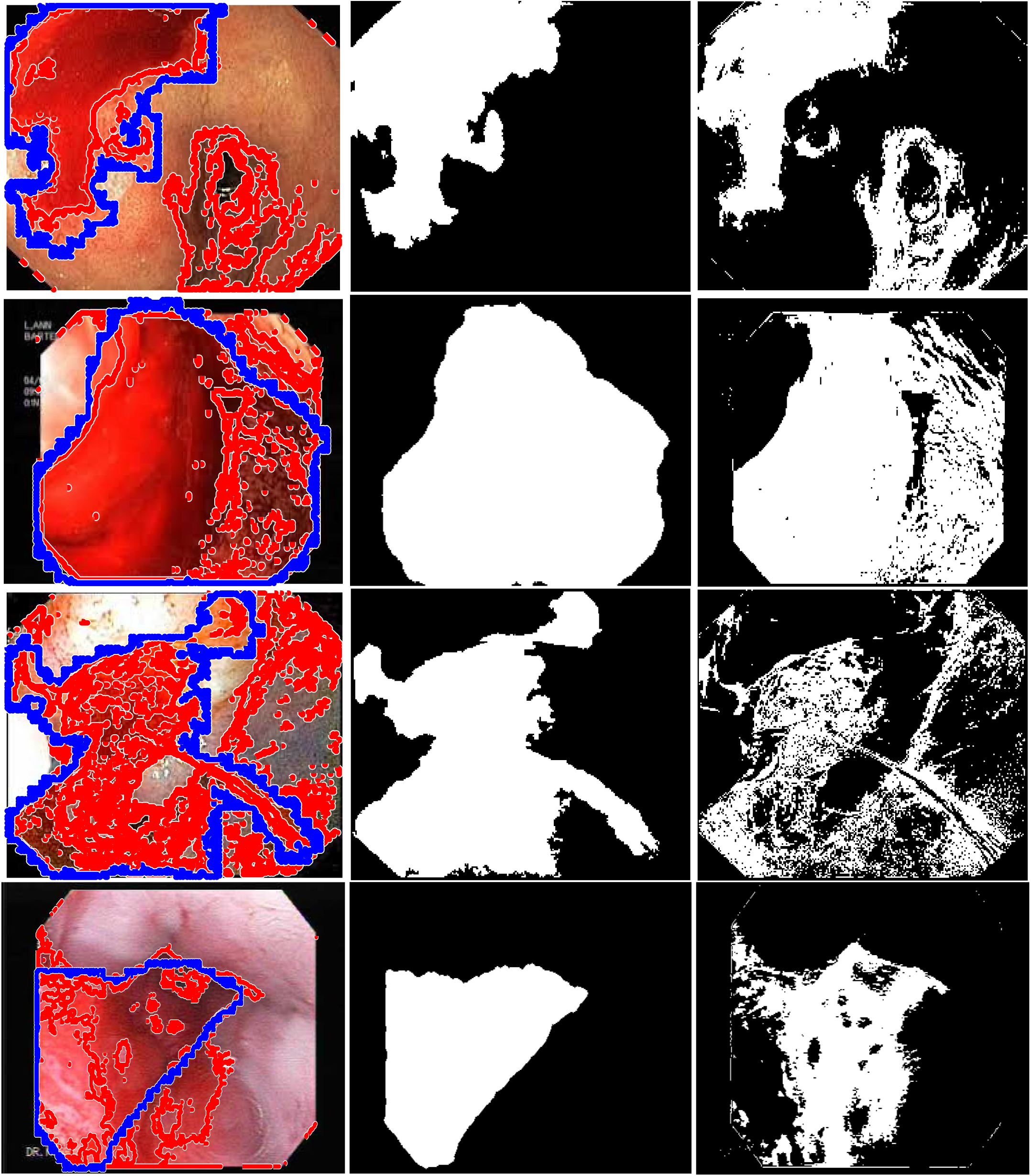
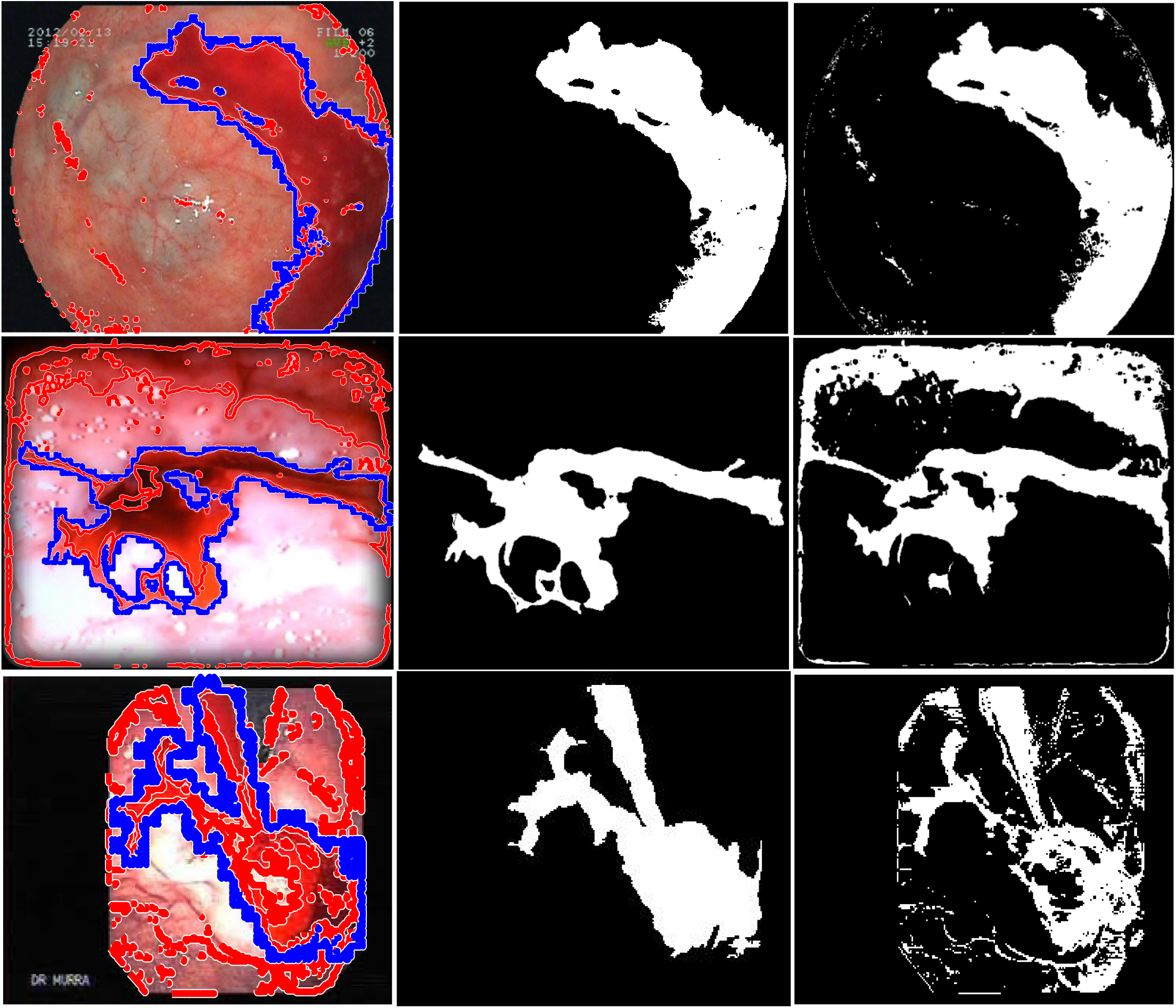
The Statistical Features of Dataset Used. lesion boundaries from the entire endoscopy images (a) Fuzzy connected object (b)Region of interest (ROI) (Ground truth of Lesion Segmentation) The labelled infected areas on endoscopy images (c) Visualization of a fuzzy segmented image.

Ground truth (Mask) generation: the initial phase of the object segmentation, we used the official dataset released at GitHub without changing any settings. Clinical image capture (input) from the study finds region of interest enclosing object. GT label was provided per image. It is shown as the Figure 2(b). Binary Stack a stack of binary images is outputted, each of those representing a cluster: a white pixel belongs to the current slice cluster, a black not. It may be useful to extract back the original region.

Analysis 2D pixel space information: This DuS-KFCM segmentation approach was applied to the colocalization image created as a multiplication of the 3 RGB image channels and GLCM feature metrices.

The extracted GLCM and statistical features were serially fused. The correlation-based feature selection method chose the informative features. The resultant optimal feature vectors were fed to the clustering method (DuS-KFCM) for the fuzzy stack.

The output of these three steps is finally merged into the 2 clusters. Segmented images from a novel feature descriptor put into the refinement and segmentation network for image high level feature learning: Figure 2(c) visually Identified lesion area. This improvement is further compared with the seed template image to obtain the image segmentation evaluation result. Performance comparison of GMM, FKM, FCM and proposed method (Deep DuS-KFCM & Dus-KFCM) for GB images based on cluster indices. To check the further robustness of our method, we used our proposed algorithms trained on GRS dataset. The performance of the method was evaluated using the benchmark dataset: Gastric Bleeding dataset and GRS dataset.

## 5. Experiment #1: Introduction of DuS-KFCM

Partitioning of an image into several constituent components is called image segmentation. The SKFCM algorithm introduces a kernel function and spatial constraint based on fuzzy c-means clustering (FCM) algorithm, which can reduce the effect of noise and improve the clustering ability. To address these issues, we propose a novel deep fuzzy DuS-KFCM for GB pixelwise classification. Integrating the colour and texture is more effective than the texture features using GLCM to represent images.

Feature plays an important role in the processing of medical images. The different features of an image include color, texture, shape or domain specific features. Texture is considered as one of the important features of an image. GLCM texture features have been widely used to characterize biomedical images. The proposed method is used to find GLCM and Fuzzy Clustering for GB Features Classification. The goal of the proposed method is to output useful information for cancer boundary through medical image segmentation and be efficient in classifying cancer. This study also attempts to combine several methods to create an effective segmentation. A flowchart of this process is illustrated in Figure 3.

**Figure 3.**
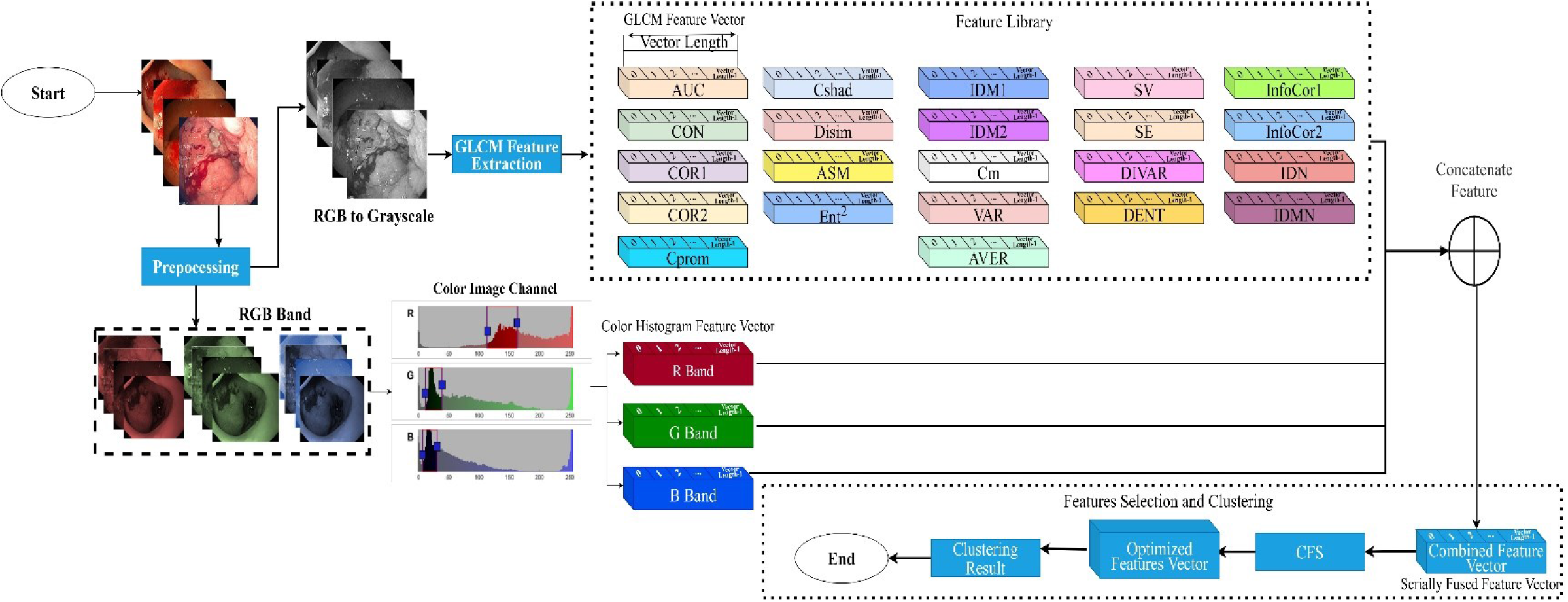
The Feature extraction system overview.

Depend on both pixel values and spatial inter-relationships, A new method for segmentation of endoscopy images is presented based on statistics of gray level co-occurrence matrix (GLCM) and the spatial kernel fuzzy C-Means clustering. The experimental results demonstrate its utility in the classification of various gastric tissue features. Subsequently, the Gray Level Co-occurrence Matrix (GLCM) features and 22 statistical features including energy, contrast, entropy, correlation and homogeneity were computed for each visible band (R, G, B). The computed features were serially fused and the best features (those that were optimally discriminatory) selected using a deeplabv3+ with resnet50 in Experiment #2 for clustering. The GLCM of an image of size 13*22 is a matrix G---Spatial Interdependence Matrix (SIM).

the primary motive of developing a feature descriptor is to efficiently capture both the color and texture information present in the image. This makes the feature a multipurpose descriptor that can work for a large variety of images belonging to different databases.

Gray-Level Co-occurrence Matrix (GLCM) methods is one of the well-known texture analysis methods to estimate the image properties related to second-order statistics [4]. The matrix is designed to measure the spatial relationship between pixels. This method is based on the belief that the information of texture is resulted from some relationships. Each entry (*i,j*) in GLCM is in accordance with the number of occurrences of a pair of gray levels *i* and *j*, which are separated by distance d as a distance in the original image.

In this study, 22 features were used as it is believed that the 22 features can distinguish objects well [5]. These features were as follows:

The gray level co-occurrence matrix (GLCM) is a classic spatial and textural feature extraction method. The Gray Level Occurrence Matrix (GLCM) is used to extract the texture features for getting the optimal threshold. GLCM Features [6]. After obtaining the segmented images form the above subsection, we extract GLCM features and store them. GLCM stands for Gray-Level Co-occurence Matrix. Texture refers to visual patterns or spatial arrangement of pixels that regional intensity or color alone cannot sufficiently describe. Texture Analysis Using the Gray-Level Co-Occurrence Matrix (GLCM) is a statistical method of examining texture that considers the spatial relationship of pixels. At the output of these algorithms, a separate feature matrix is obtained for each image. The GLCM algorithm generates a feature matrix with 22 image feature and 1 class information for each image. GLCM features when extracted in various color space show better representation of human perception compared to using RGB color space. Each element of a GLCM matrix provides too “microscopic” a view of a texture in an image. What we need is a larger “macroscopic” characterization of the texture from the information contained in a GLCM matrix.

The application used the spatial gray level co-occurrence matrix (GLCM) to extract texture features. Fuzzy clustering analysis transformed images to the hybrid Spatial feature vector.

In addition, a feature selection procedure using information theory has been applied to identify and select the best features that provide useful information about the characterization of the GB. For each segmented image, Normalization of feature values is necessary preprocessing step before training classifiers. The algorithm is enhanced by normalization evolution.

In any process which requires exploring data, it is important to perform a preprocessing of them. One of the most basic methods is the normalization, which is especially useful when the features have different units and scales. There are two widely known procedures to make a linear scaling of the data between the interval [0, 1] (it is also common [-1, 1]). Let *x* be a particular feature value and let *x* min and *x* max be the respective minimum and maximum values of that feature. Then the max-min scaling feature normalization is defined as follows:

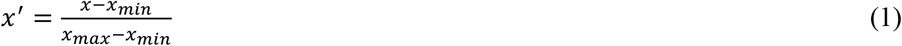

where *x*′ is the normalized feature value.

This stage is necessary to remove redundancy from the input data which only contributes to the computational complexity of the network without providing any significant improvements in the result. Not all features are correlated with disease. Must include only relevant features to avoid misclassification. When many features are extracted, the complexity of the problem description becomes high, making difficult to build a good classification system. Feature selection defines a topic commonly used in data mining to select the more significant features. Information theoretic feature selection can generate features importance of input features, which can be used to analyze features’ effectiveness. The increasing dimension of the dataset makes the classification more difficult and the development to a certain extent can cause curse of dimensionality. For high-dimensional data, firstly, dimensionality reduction is carried out, and then data after dimensionality reduction is input into the learning system. Then, correlation-based feature selection (CFS) segmentation parameter selection dimensionality reduction methods are used to compare the effect of protein GB localization overall prediction accuracy by using these dimensionality reduction methods. segmentation results of GB lesion of the gastric dataset by selecting different dimensionality reduction methods and different dimensions choosing different dimensionality reduction methods and dimensions have a significant effect on the accuracy of GB segmentation.

These two feature descriptors, color histogram and 22 second order statistical texture feature, are concatenated in order to utilize both the global and local information of the image respectively which found to be beneficial in our experiments. Moreover, we calculate the GLCM of the feature map obtained rather than computing the histogram to maintain the information of the spatial correlation. A feature vector is a method to represent an image or part of an image. A feature vector is an n-dimensional vector. Effects of selecting 3 different dimensionality reduction methods and different dimensions on the overall segmentation results of GB lesion location in gastric dataset. As the user selected segmentation features vary, the resulting objects will change. vectors data preparation and clustering.

As shown in the Figure 7. we choose the following three feature evaluation algorithms. The search for effective features in a feature-rich area is performed. If the *p* value is greater than the critical value at 0.05 significant level, then the feature is significant, and it is selected. We select 8 of the best features from the feature library including F2, F6, F7, F17, F18, F19, F21 and F22. The data sets are listed in Table 2.

**Figure 4.**
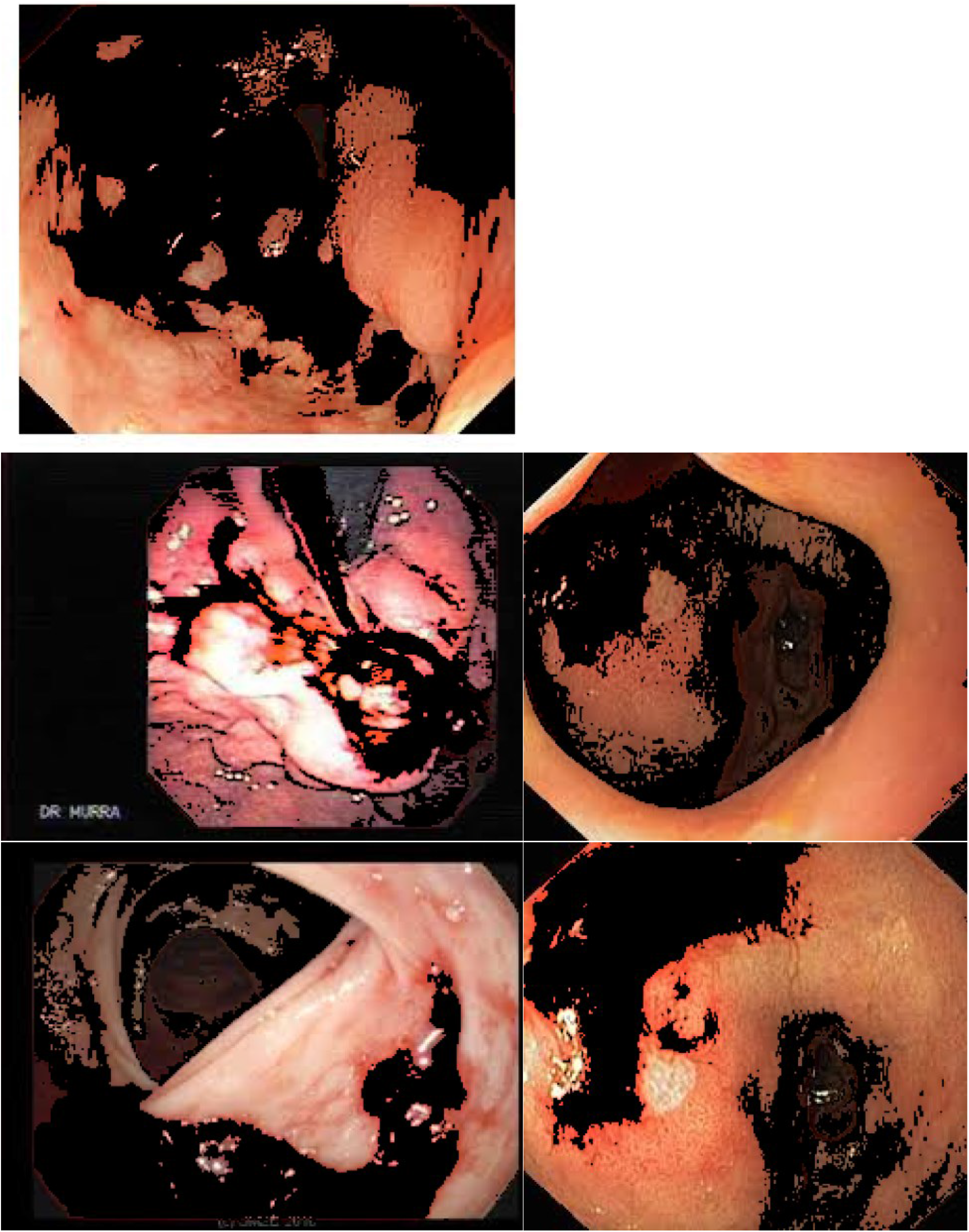
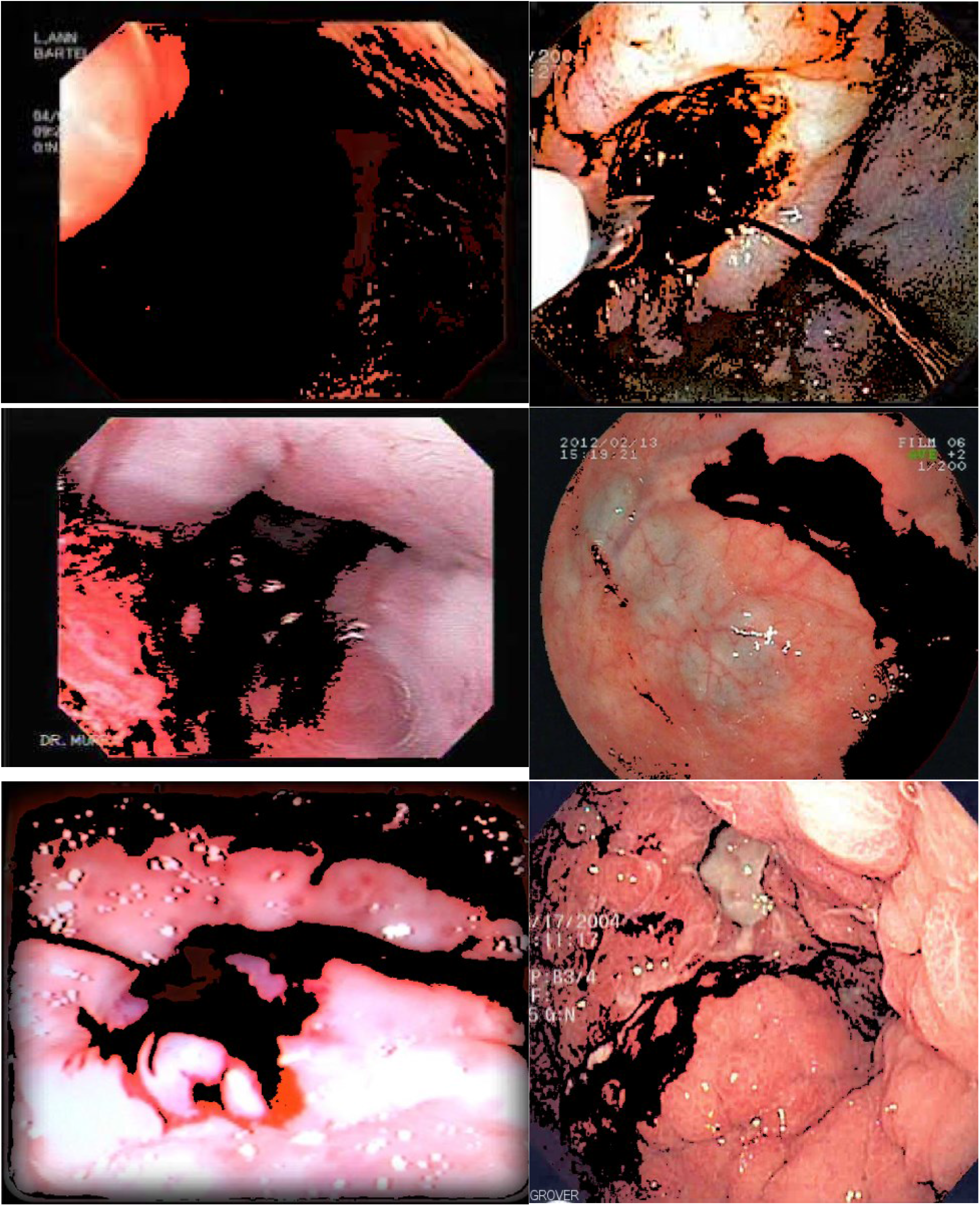
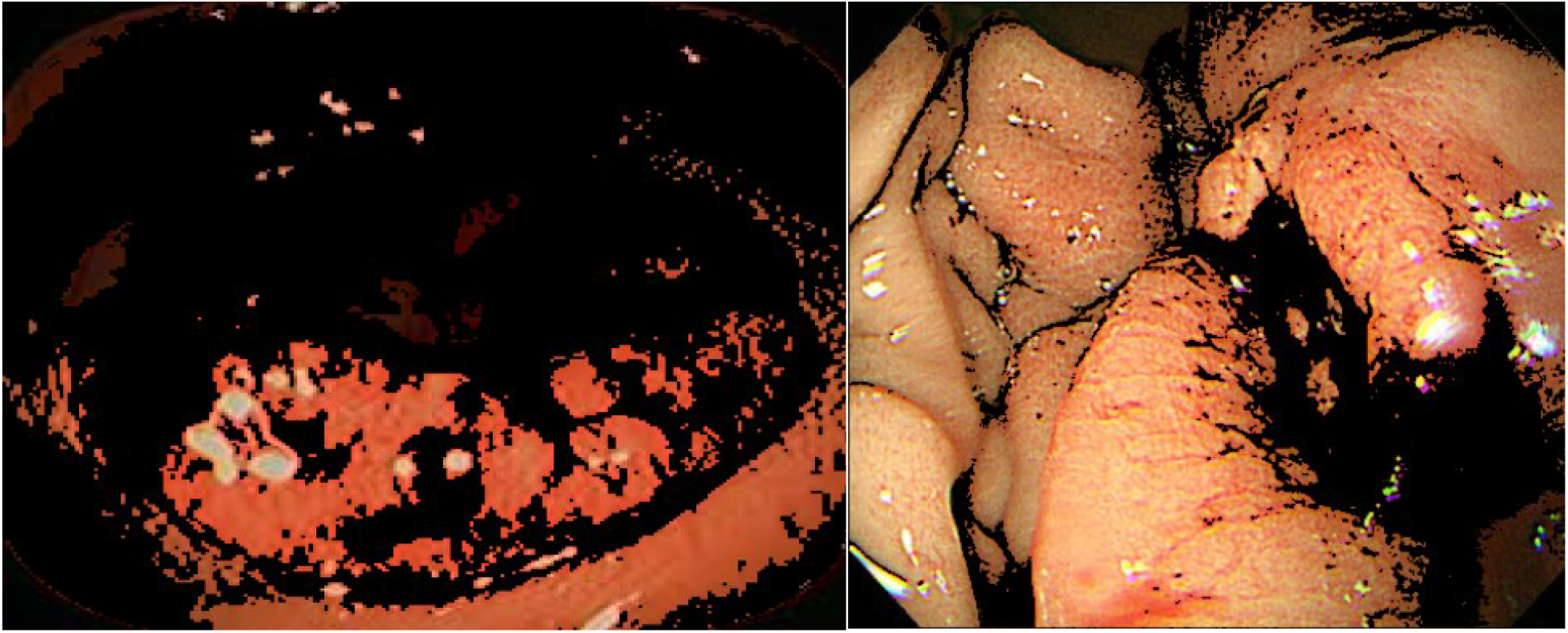
GLCM-based features on normal Area

**Figure 5.**
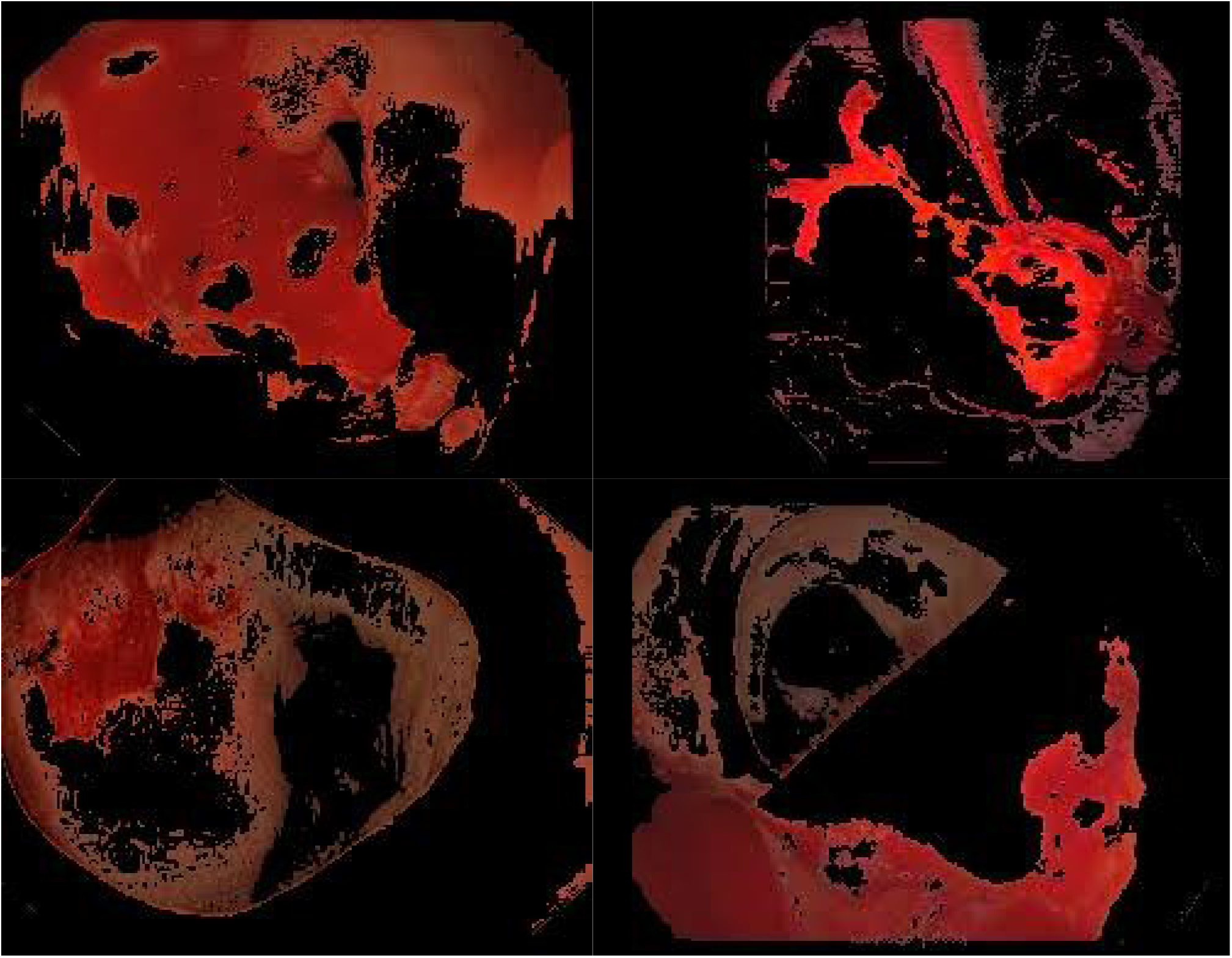
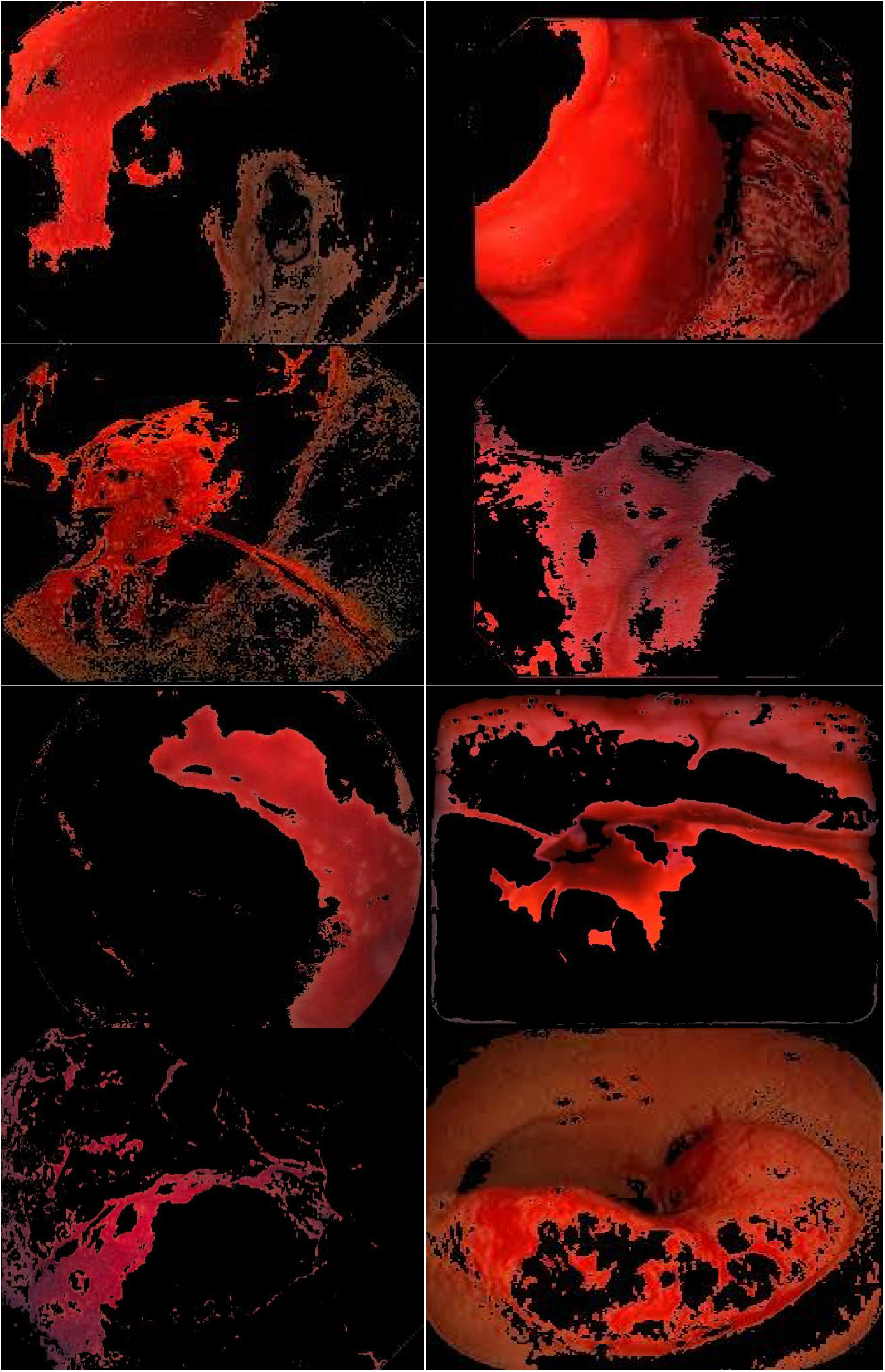
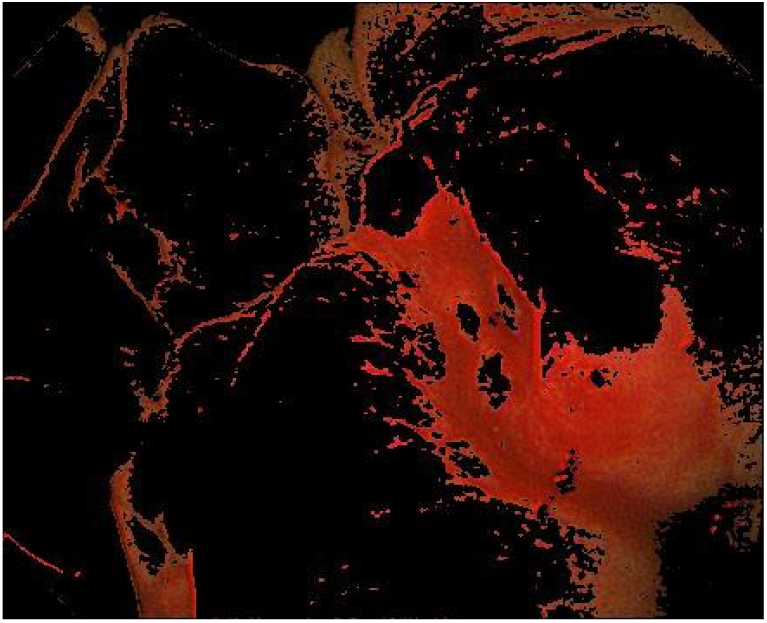
GLCM-based features on GB lesion areas

**Figure 6.**
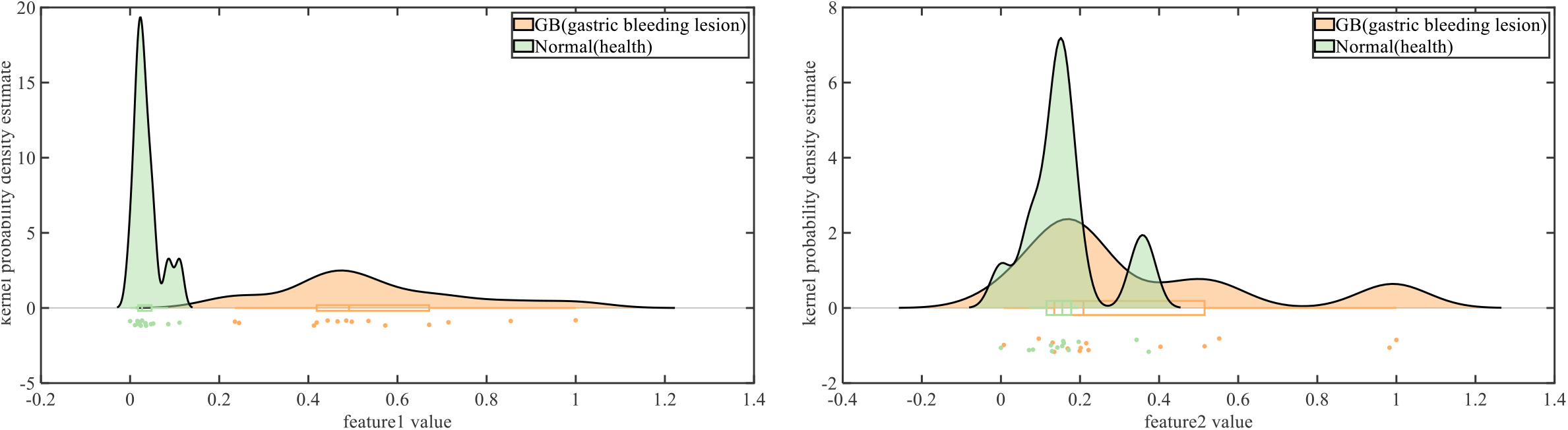
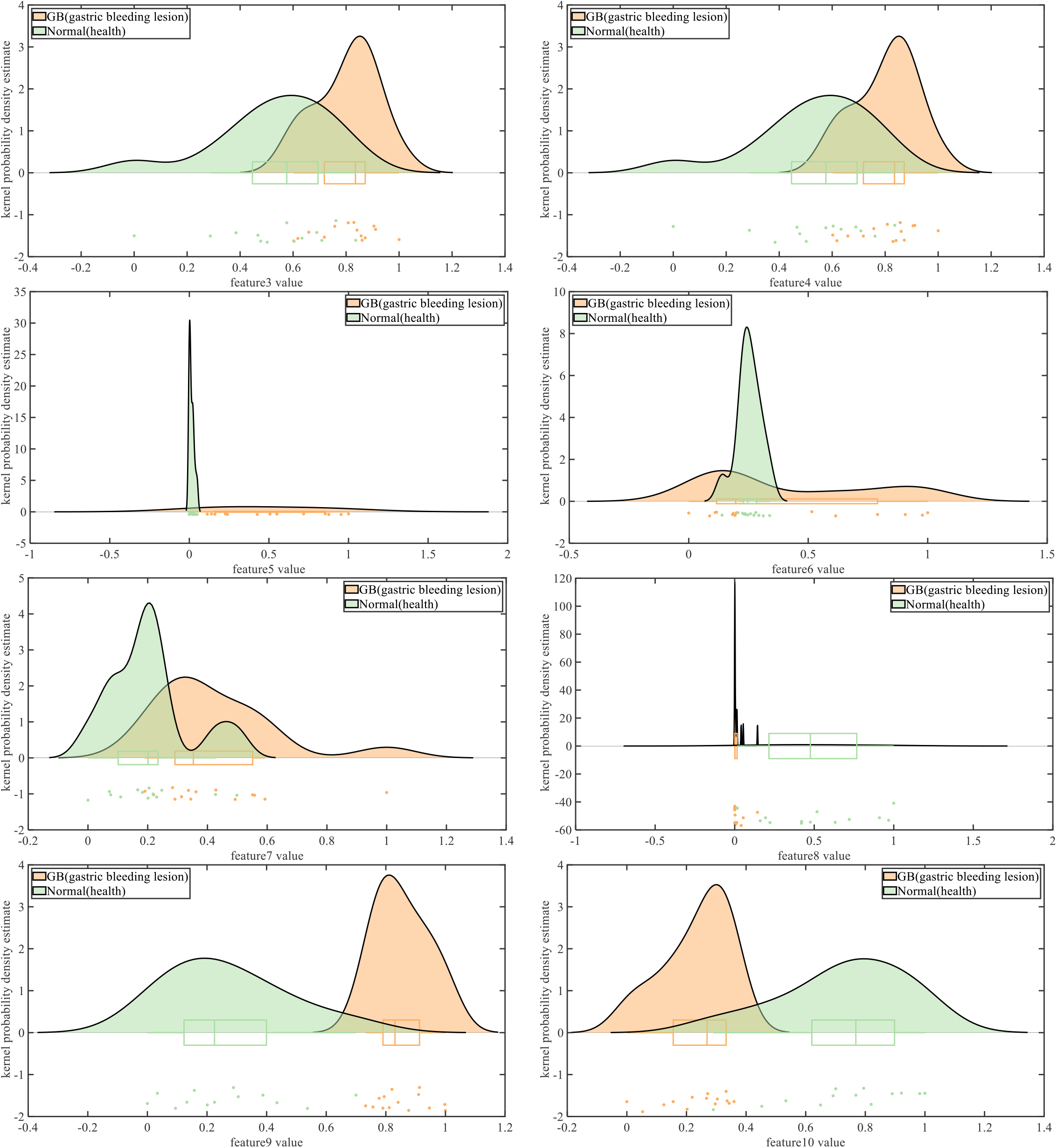
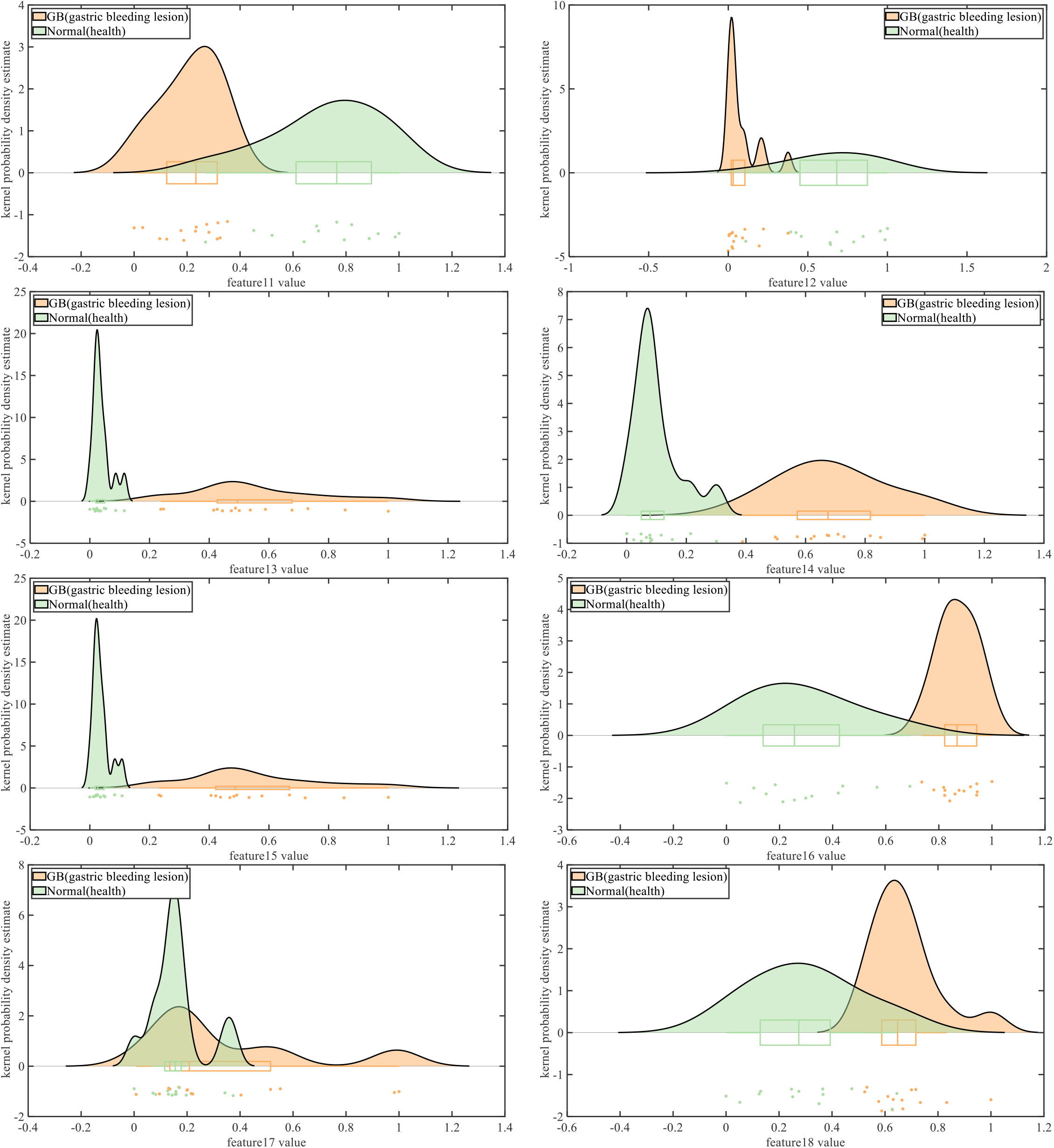
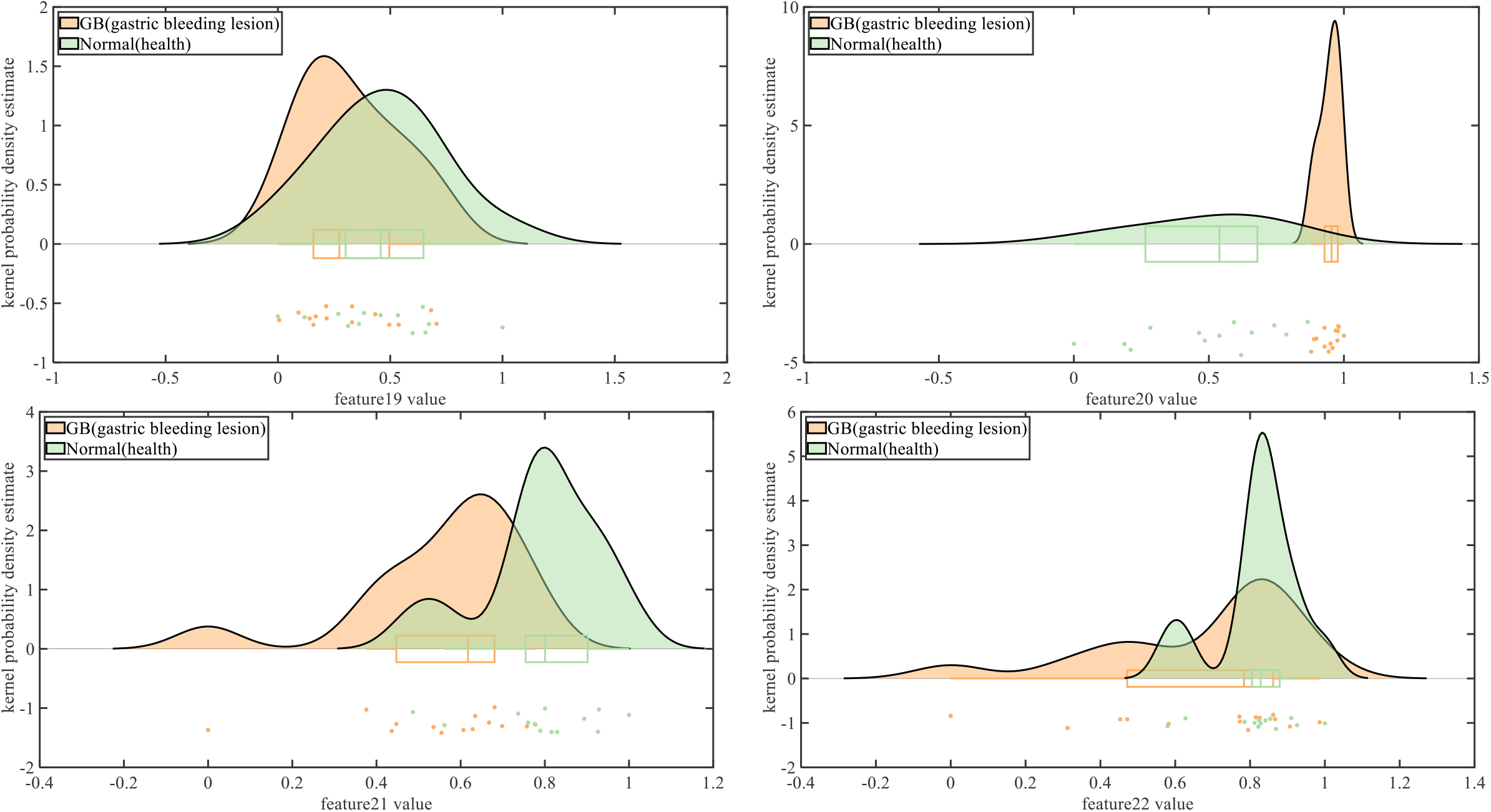
GLCM-based features exploration using the raincloud plots. (Statistical significance of the gastric bleeding clinical information in the GLCM.)

**Figure 7.**
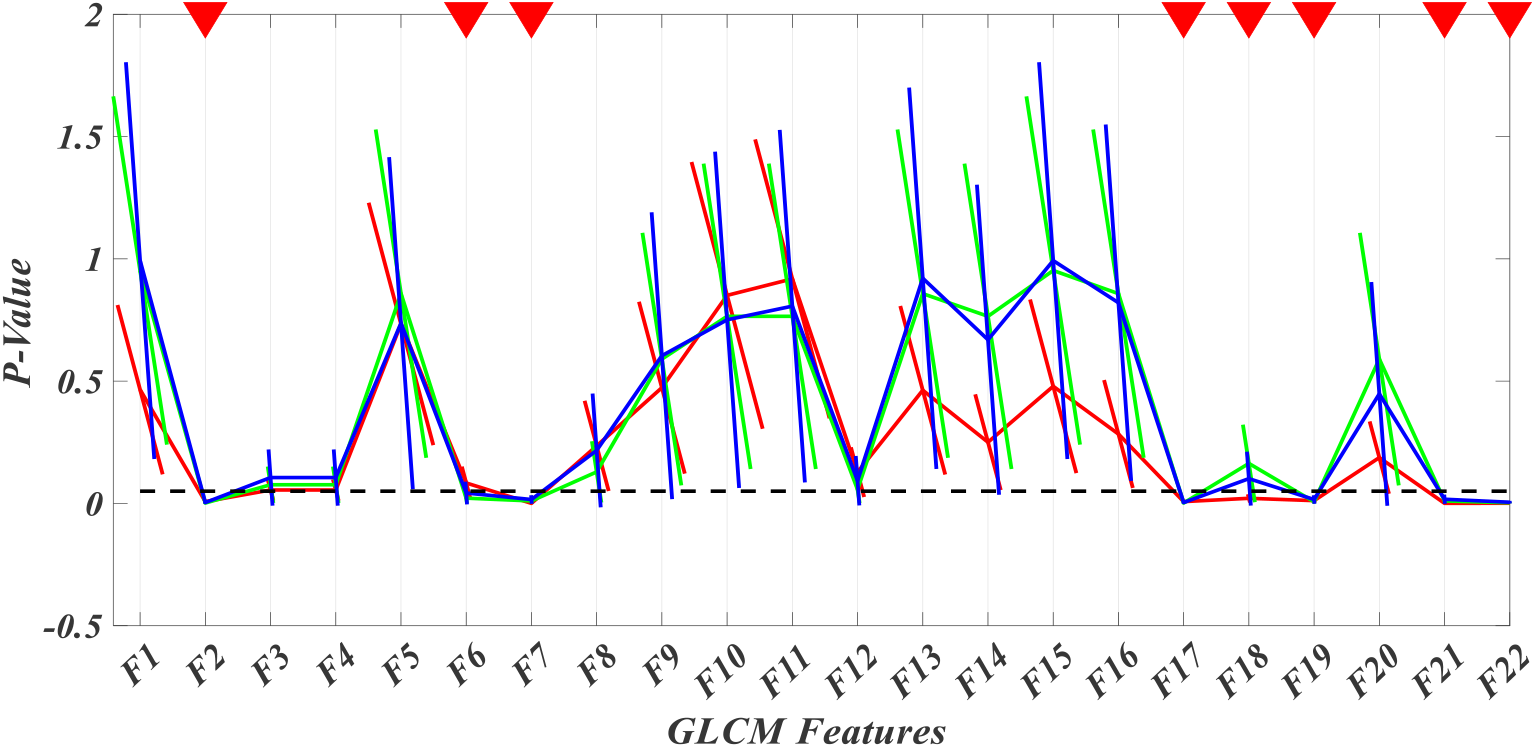
Feature selection Operations based on 3 method; (Black baseline: specifying a scalar value for significant difference(p-value); Red curve: Errorbar diag of Pearson Correlation-Based Feature Selection; Green curve: Errorbar diag of Kendall Correlation-Based Feature Selection; Blue curve: Errorbar diag of Spearman Correlation-Based Feature Selection;

**Table 1.**
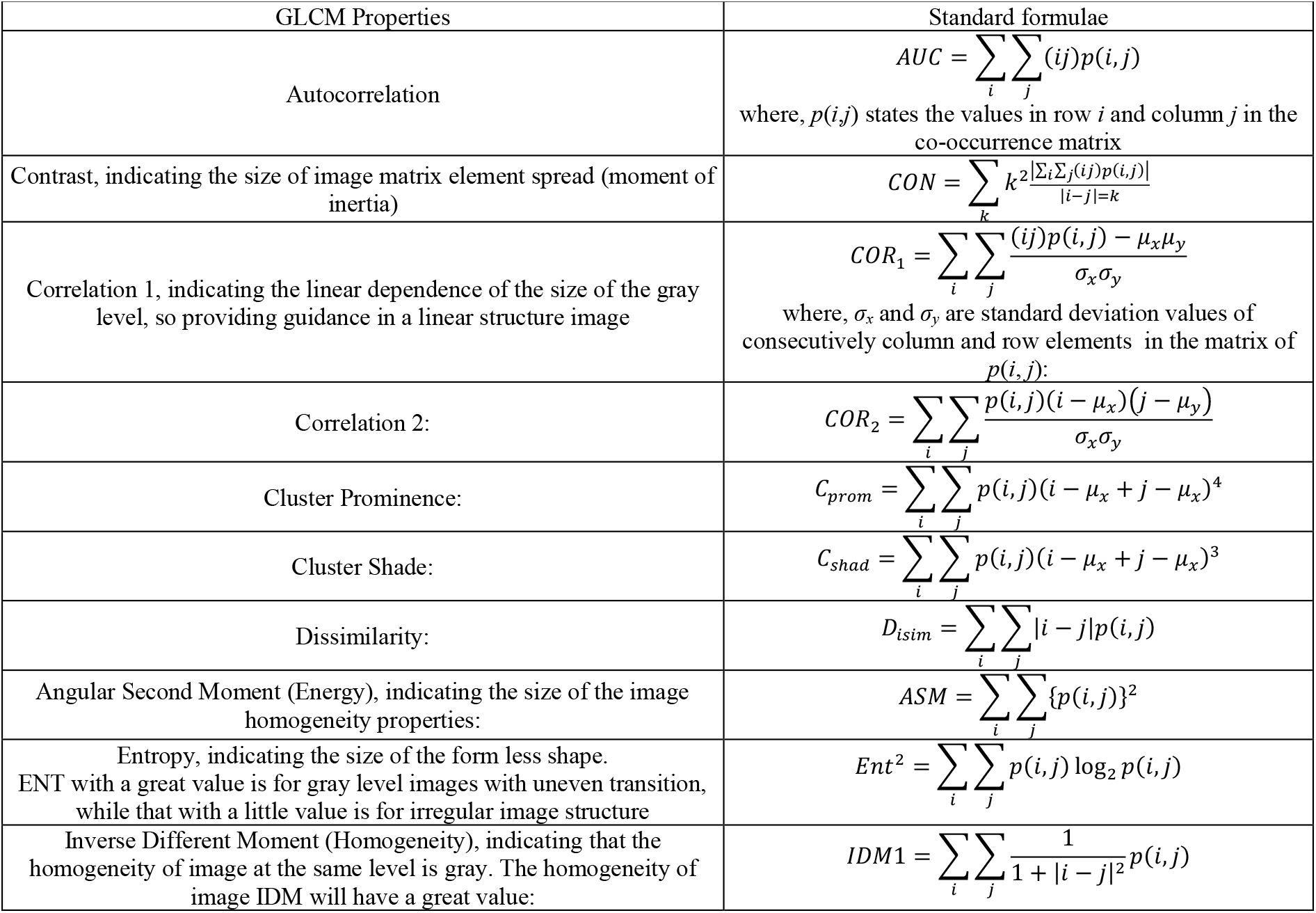
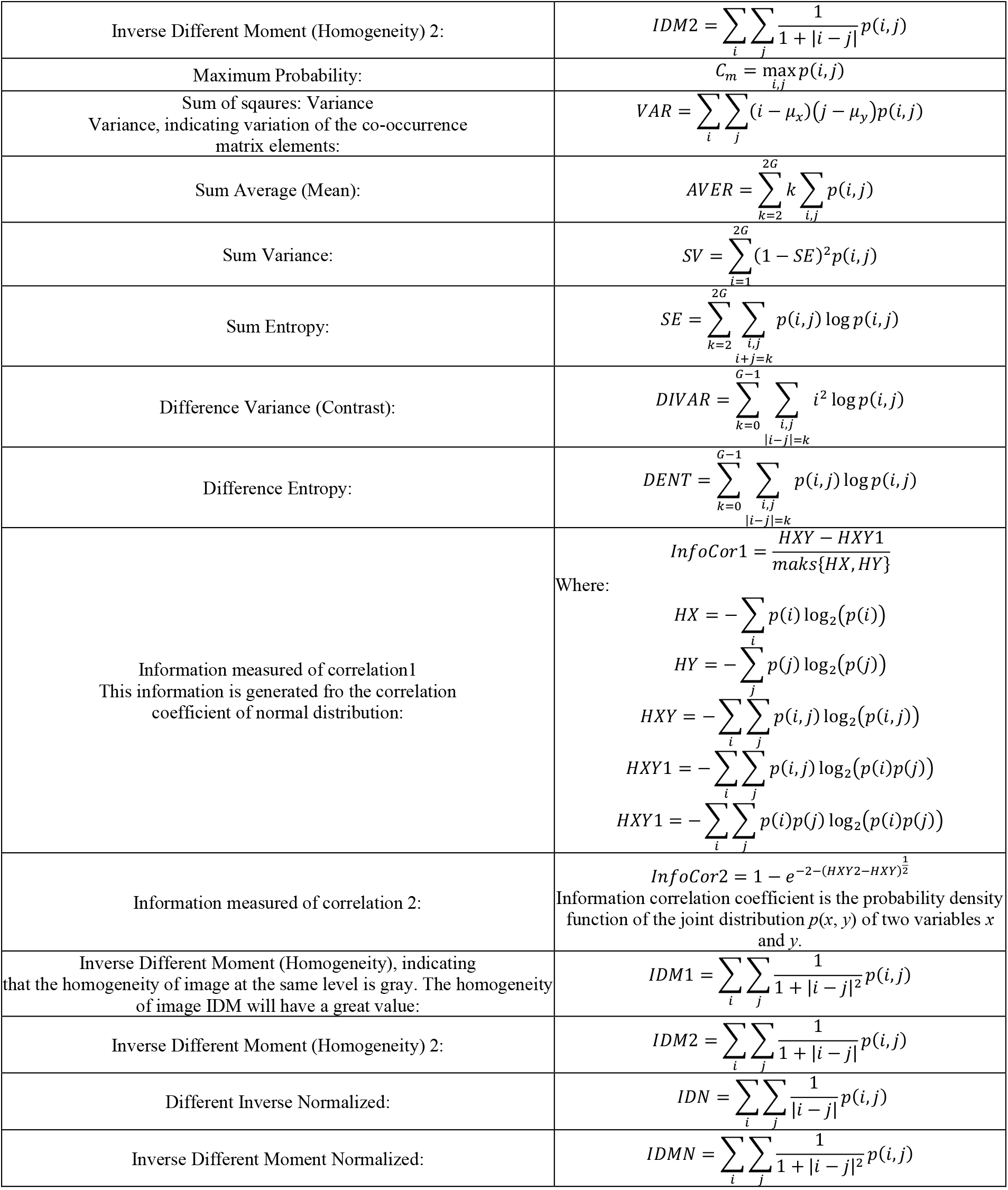
GLCM properties and corresponding formulae

**Table 2.** Features Selection(P-value) Dimension Reduction (Selection of dimensionality reduction method and optimal dimension)

| Type of correlation | F1 | F2 | F3 | F4 | F5 | F6 | F7 | F8 | F9 | F10 | F11 |
| --- | --- | --- | --- | --- | --- | --- | --- | --- | --- | --- | --- |
| 'Pearson' | 0.4651 | **0.0074** | 0.0548 | 0.0548 | 0.7338 | **0.0836** | **0.0003** | 0.2352 | 0.4732 | 0.8506 | 0.9183 |
| 'Kendall' | 0.9524 | **0.0016** | 0.0763 | 0.0763 | 0.8577 | **0.0216** | **0.0101** | 0.1289 | 0.5900 | 0.7650 | 0.7650 |
| 'Spearman' | 0.9928 | **0.0036** | 0.1057 | 0.1057 | 0.7370 | **0.0425** | **0.0160** | 0.2166 | 0.6039 | 0.7507 | 0.8065 |
| Type of correlation | F12 | F13 | F14 | F15 | F16 | F17 | F18 | F19 | F20 | F21 | F22 |
| 'Pearson' | 0.1278 | 0.4627 | 0.2503 | 0.4789 | 0.2838 | **0.0074** | **0.0209** | **0.0114** | 0.1874 | **0.0005** | **0.0007** |
| 'Kendall' | 0.0573 | 0.8577 | 0.7650 | 0.9524 | 0.8577 | **0.0016** | **0.1635** | **0.0067** | 0.5900 | **0.0101** | **0.0027** |
| 'Spearman' | 0.0929 | 0.9206 | 0.6693 | 0.9928 | 0.8206 | **0.0036** | **0.1013** | **0.0160** | 0.4478 | **0.0171** | **0.0048** |

In the experiment#1, the data-level fusion was implemented. the 3 colour spaces has been chosen from where the colour texture GLCM features were extracted. Proposed fused features that contains a set of values where each value represents a certain feature. Construction of feature vector can allows be used to classify an object.

Segmentation based on the controlling parameters spatial domain features provide us with condensed higher-level information regarding the image. This vector for global color histogram thresholding and extracted the bleeding region. Integrated method for a targeted region leads to more robust segmentation.

In this 1^st^ experiment, we have attempted several approaches for the segment of lesions in gastric bleeding images. fuzzy segmentation was performed using a group of similar pixels as revealed by their class labels. The prediction was performed at the pixel level; each image pixel was classified based on the corresponding class label. The results of the proposed segmentation method are compared pixel-by-pixel with the ground truth images in Figure 8. using GLCM based feature and fuzzy color thresholding extraction. The proposed method accomplishes higher classification accuracy than other methods such as fuzzy k-means clustering (FKM), Gaussian mixture model (GMM), fuzzy c-means (FCM) clustering, and dual spatial constraint into fuzzy c-means (DuS-KFCM) clustering. Comparison

**Figure 8.**
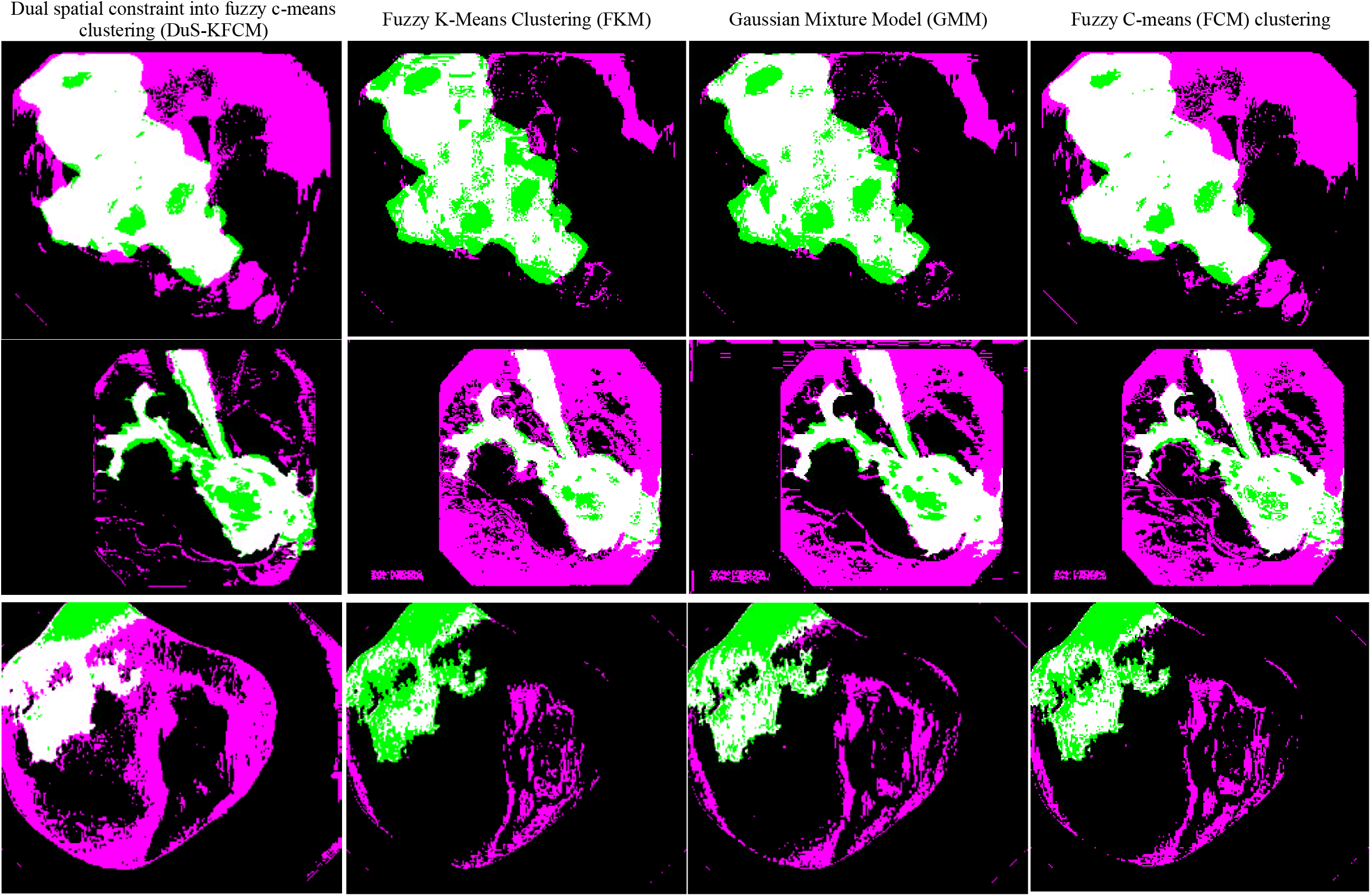
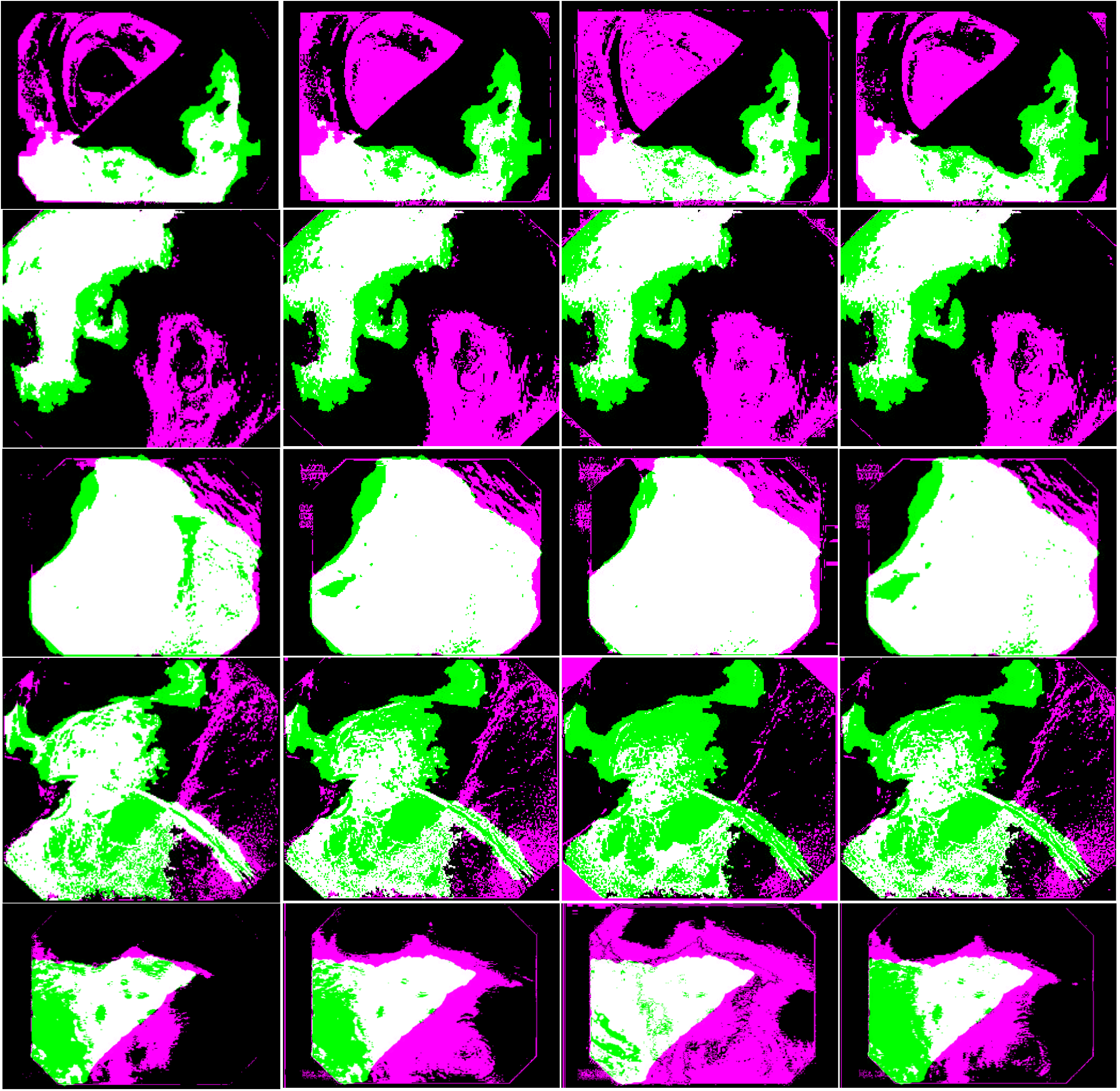
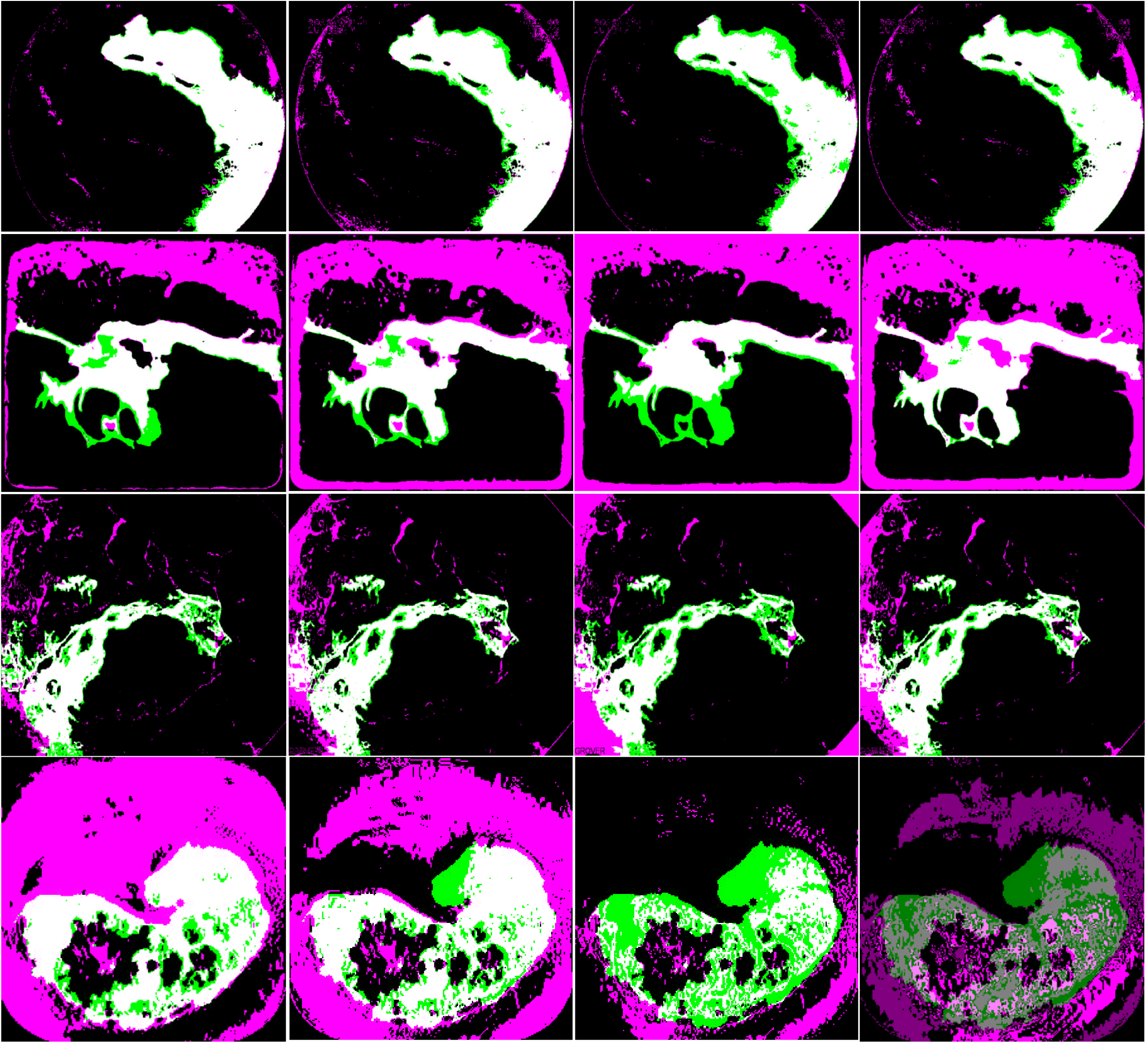
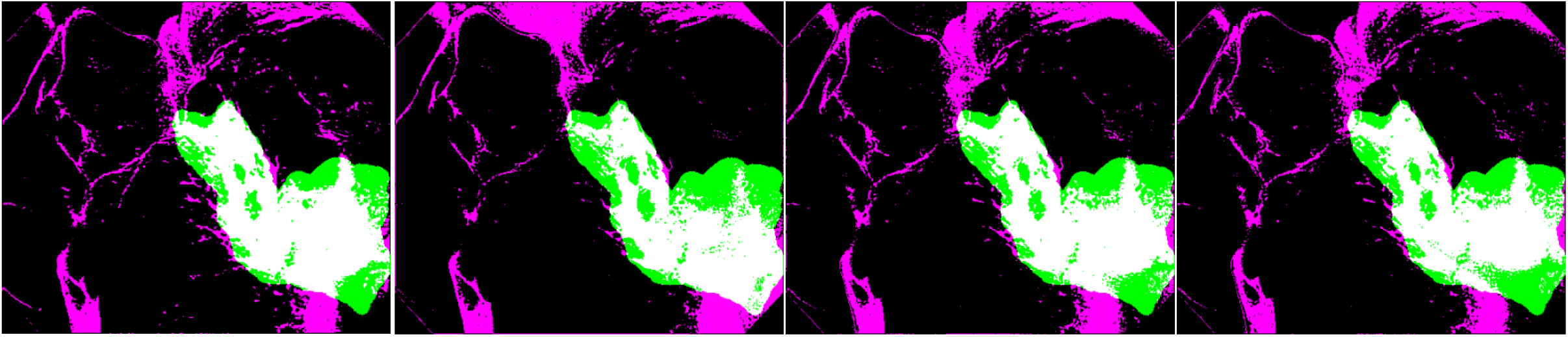
Results on the Synthetic Images: different fuzzy clustering (Proposed methods vs. Traditional threshold methods) leading to differently clustered images (Magenta area: Boundary by system; Green: Boundary by Gastroscopist; White area: fusion Image D with RA/RB as reference information)

Visual comparative analysis on GB dataset is shown as the Figure 8.

## 6. Implementation of the deep DuS-KFCM image segmentation

A significant proportion of the misclassification rate may be attributed to the small spurious regions that are frequently included in the cluster corresponding to the lesion region. To overcome the aforementioned limitations, the deep DuS-KFCM method is implemented to improve the performance of the image segmentation process. DeepLabv3 is an effective decoder that helps to refine segmentation of lesion boundaries. The refined gastric bleeding lesion segmentation based on deep learning algorithm. In our work, DeepLabv3+ with ResNet50 as backbone is used as the segmentation network.

After applying deep fuzzy enhancement to the input image set, the approach demonstrates its efficiency and effectiveness in dealing with the fuzzy segmentation. Results of performance measures segmentation based on deep DuS-KFCM for GBs proposed methods for gastric medical image are illustrated in Tables 4, respectively. The average results for Tables 3 are displayed in Table 4.

**Table 3.**
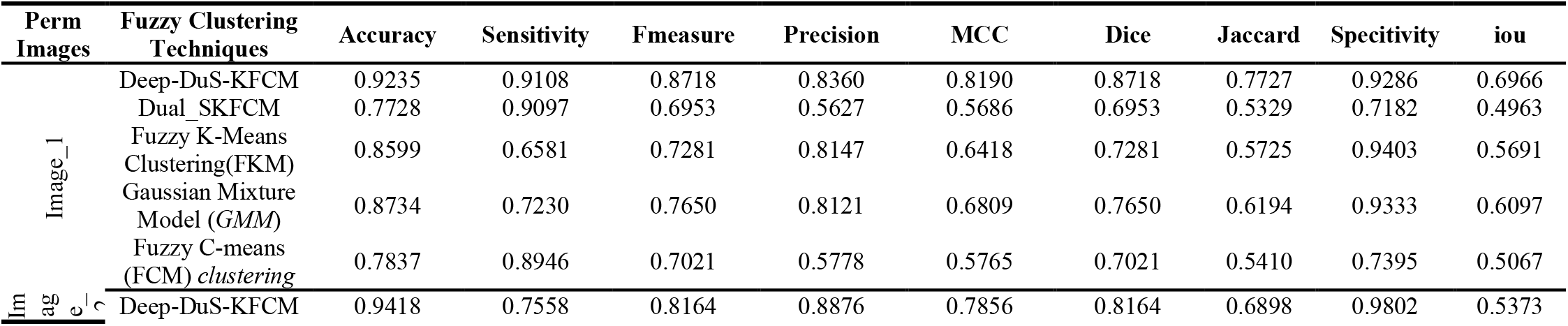
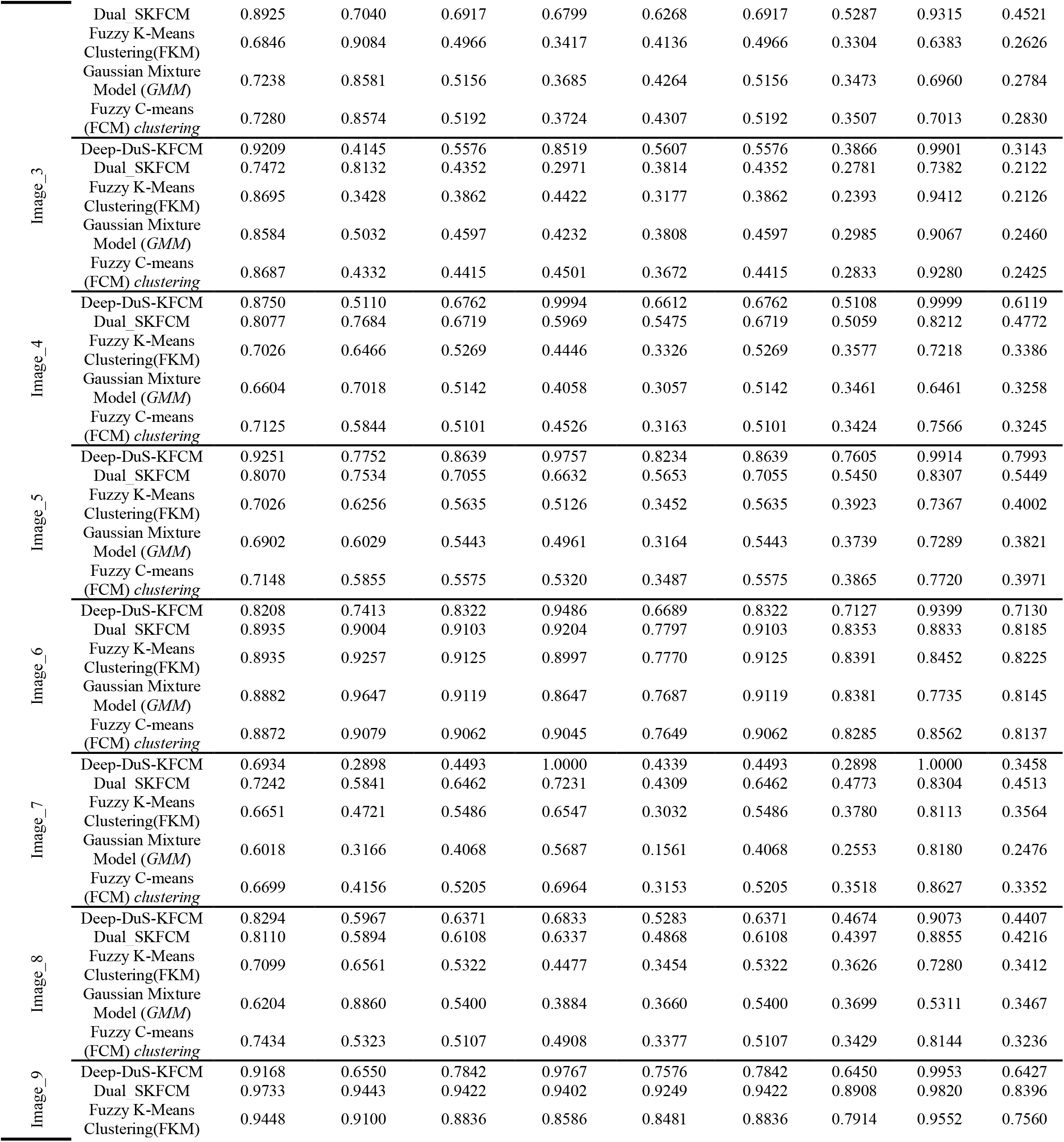
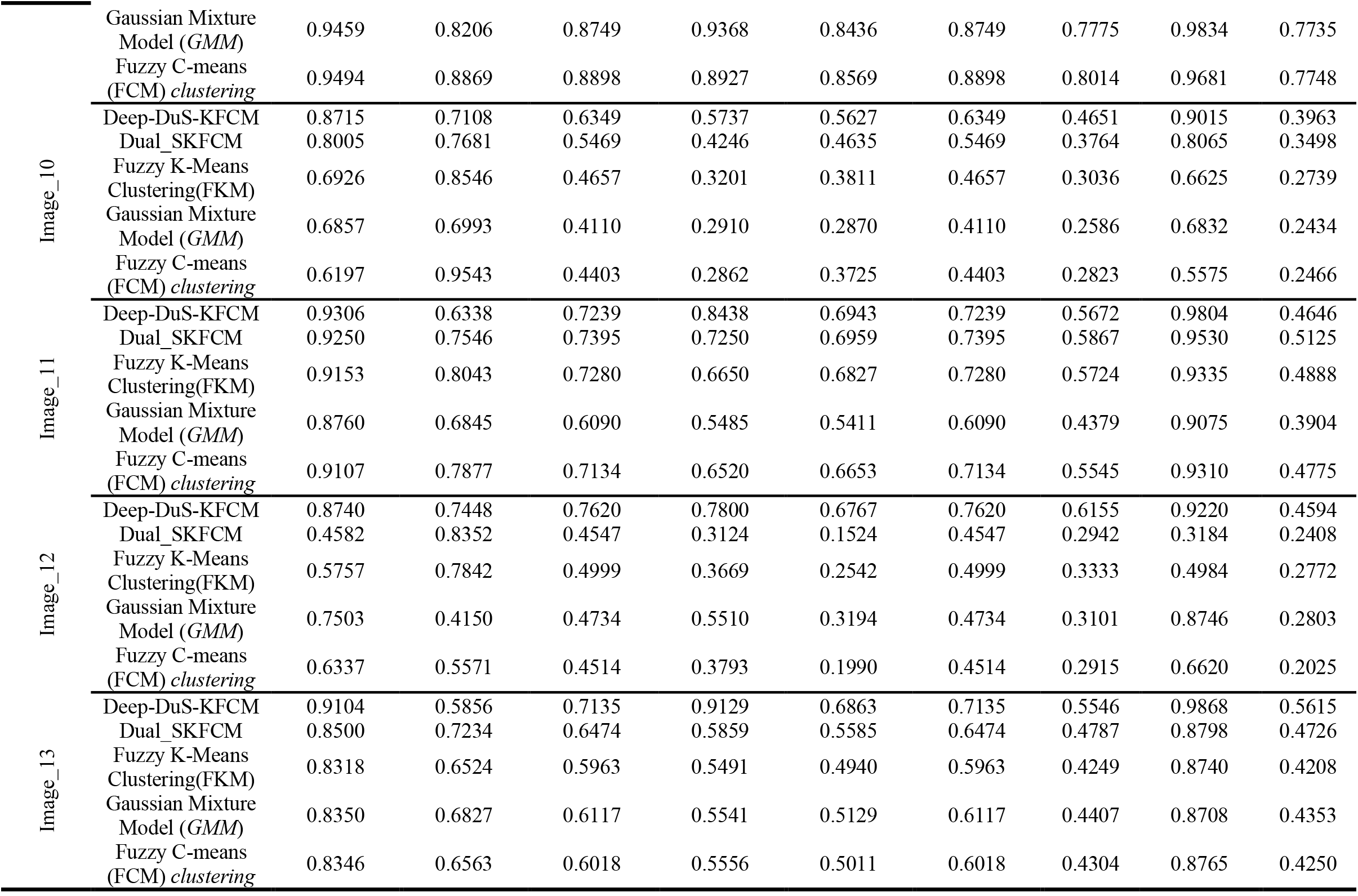
Segmentation results for a single field of view (FOV) Comparative results of the proposed method with respect to other methods.

**Table 4.**
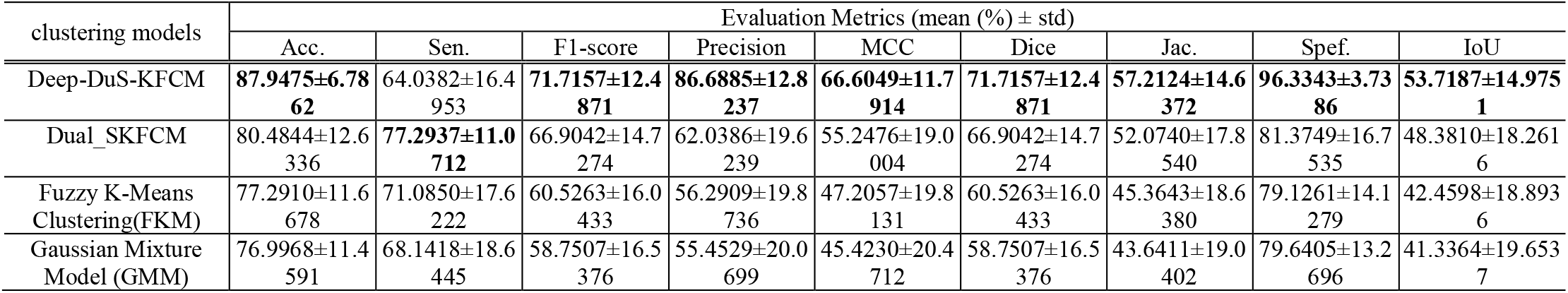
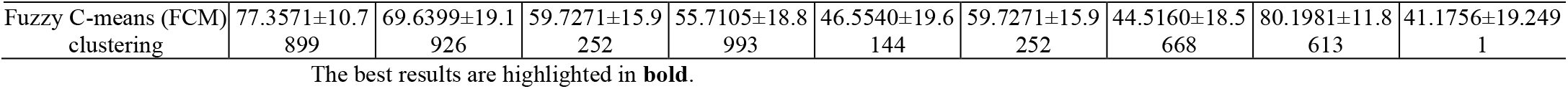
Comparison Image Segmentation Quality Scores with some related works.

Figure 9 displays the structure of the proposed system.

**Figure 9.**
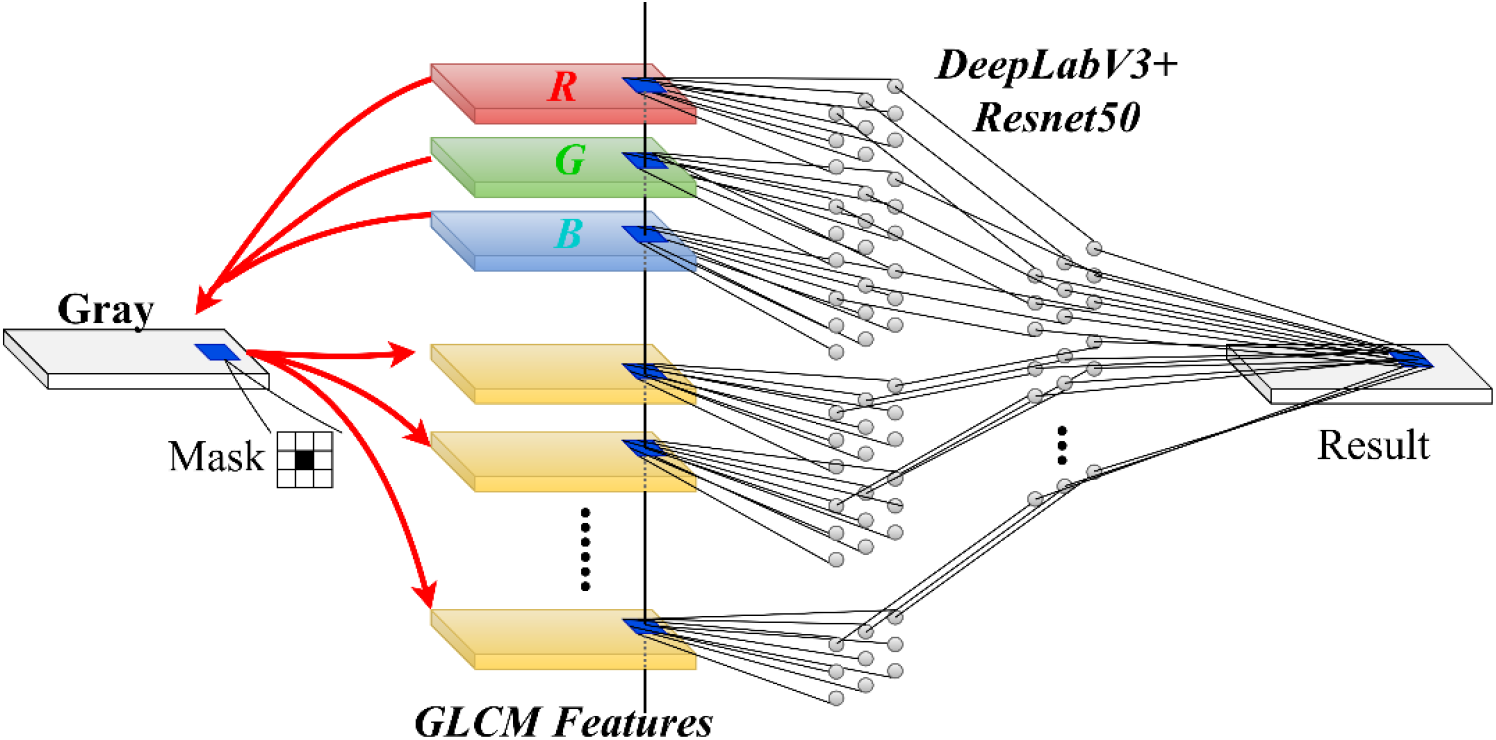
General view of the network architectures: DLv3+ with Resnet50

A block diagram representation of a neural network is shown in Figure 10.

**Figure 10.**
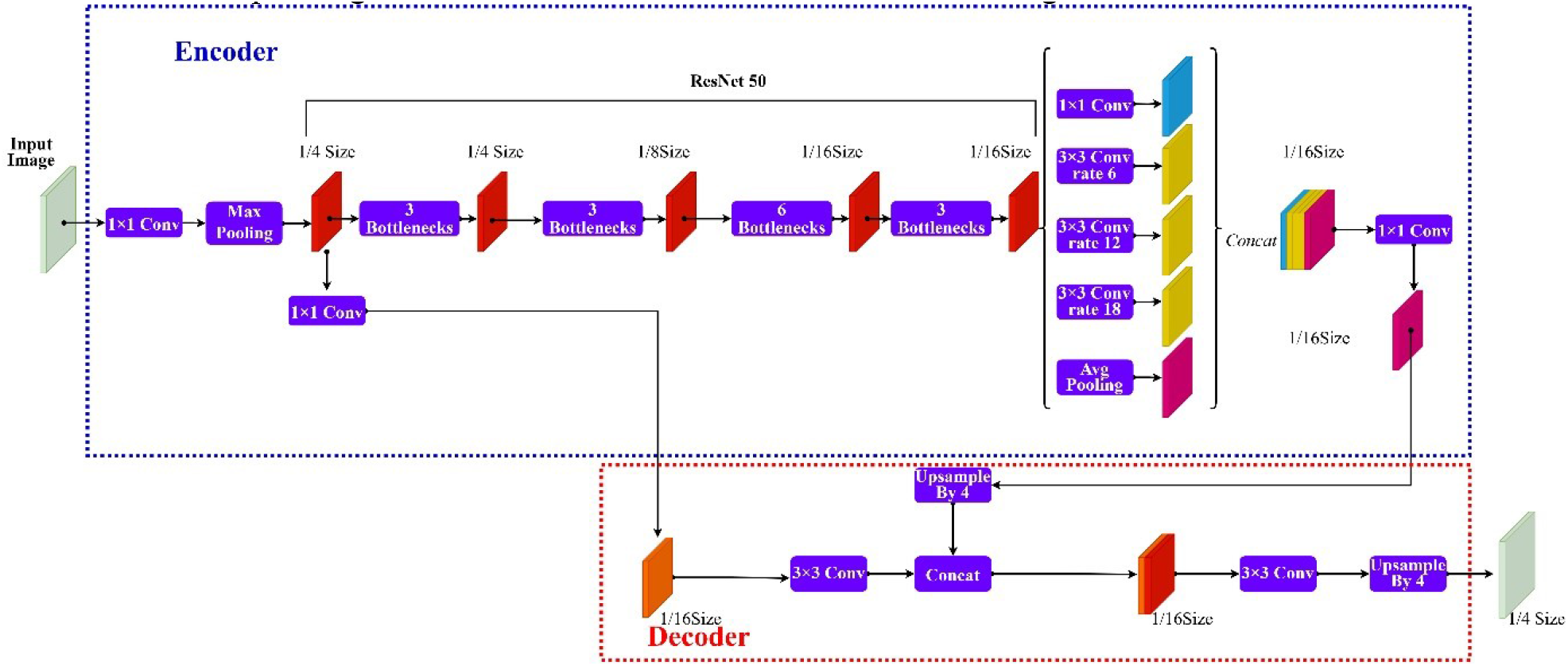
Block Diagram Representation of Neural Network

Incorporating networks into information to refine the lesion segmentation results.

The propose a novel algorithm above in which a combination of color, texture, and shape features is used to form the feature vector. The feature vector that combines all features is used to cluster images into different clusters. The ensemble (optimized) enhancement of DuS-KFCM Optimization for Medical Image Segmentation calculated the extracted GB trained image region Then, we extract high level features on the segmented image. to improve the segmentation.

The proposed segmentation outcomes are illustrated in Figure 11.

**Figure 11.**
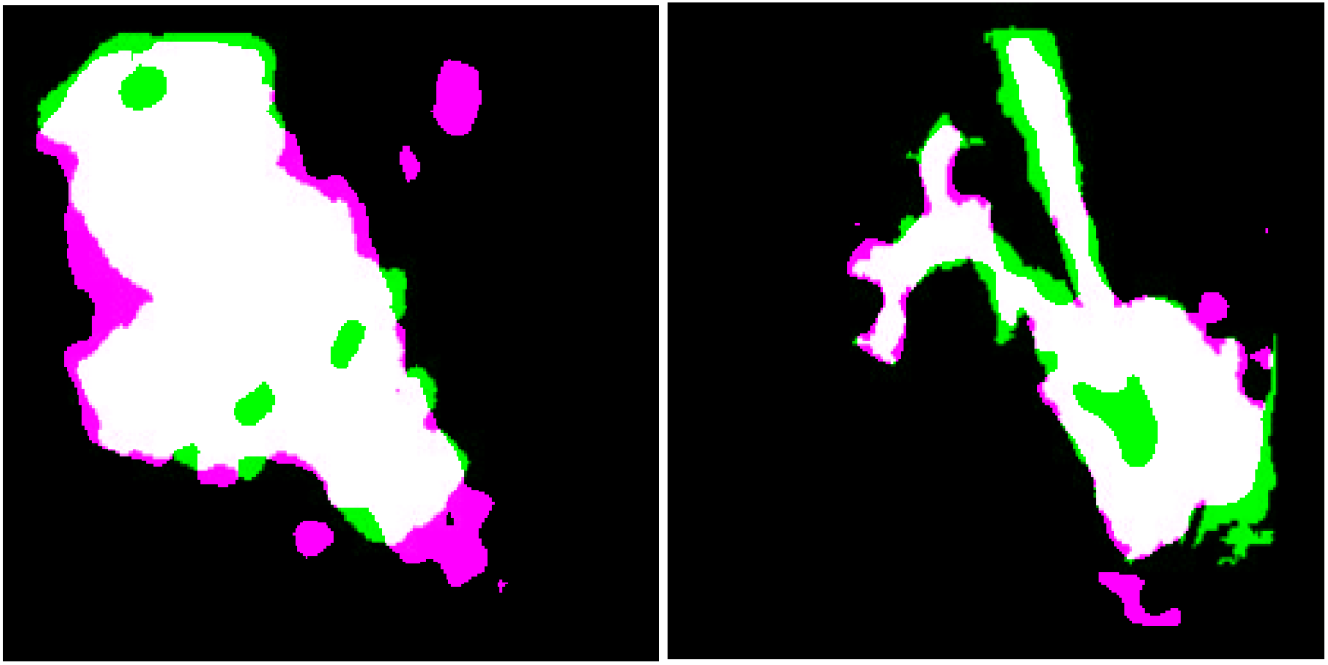
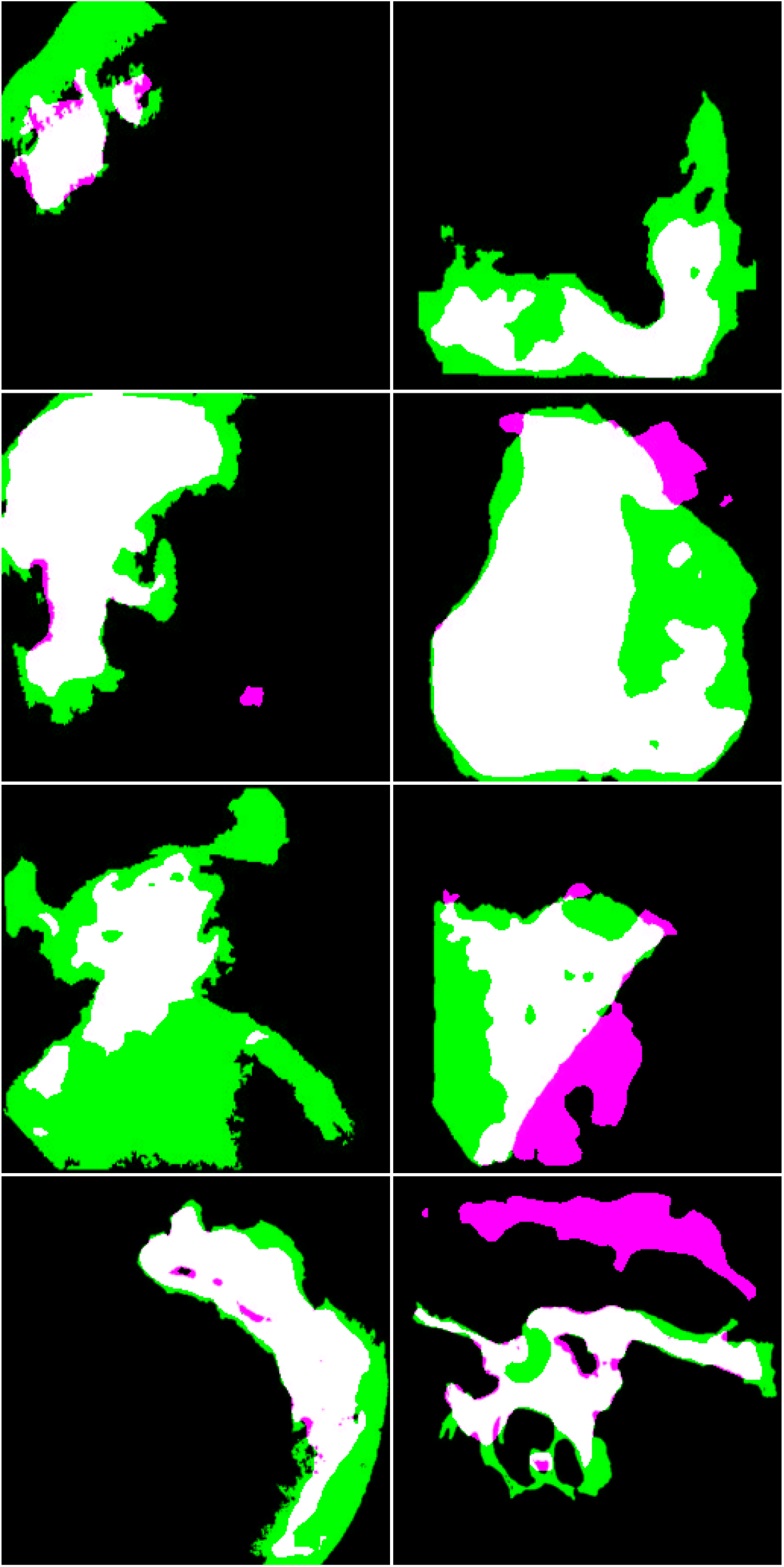
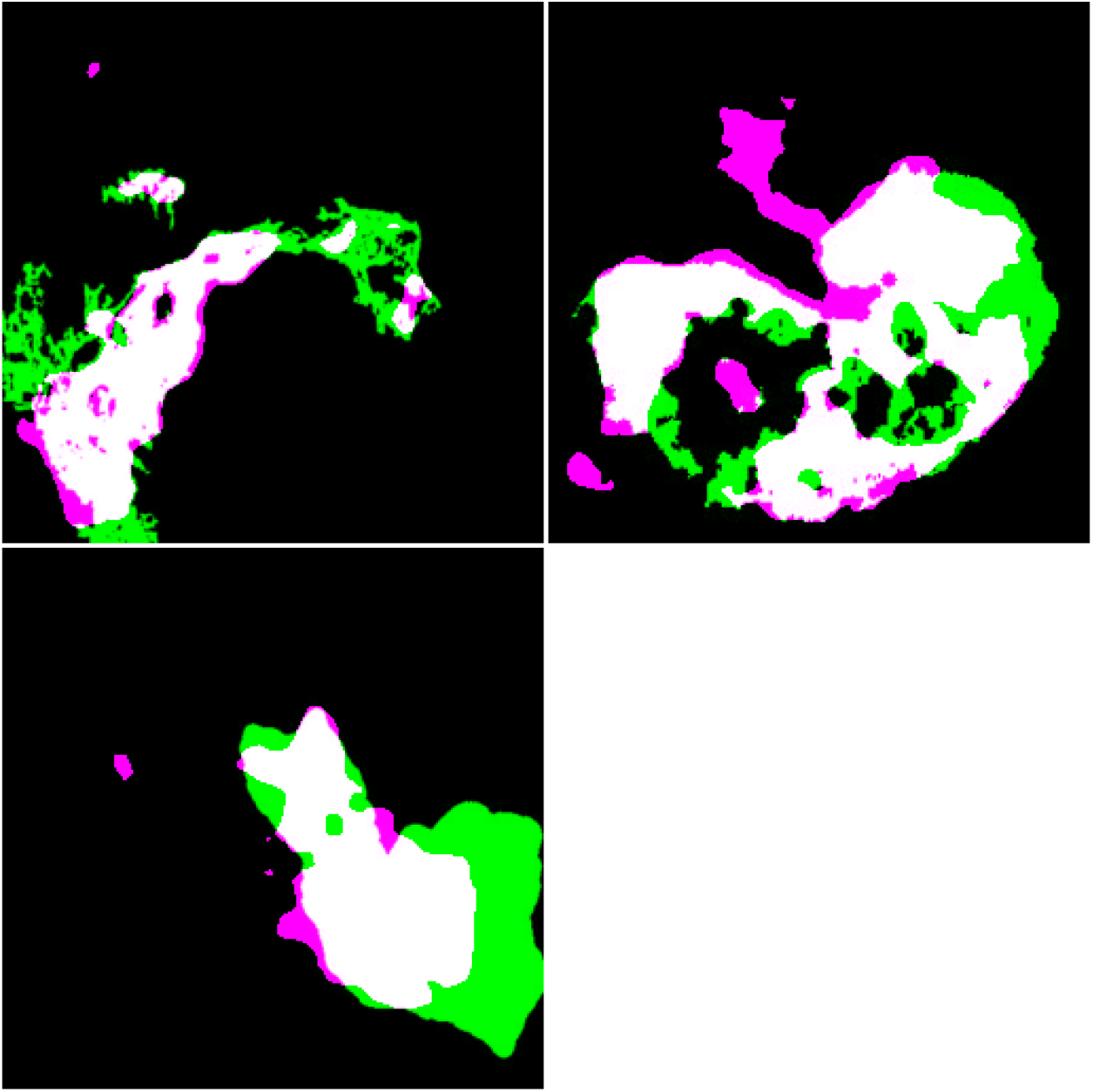
Experiment #2 Results on the Synthetic Images of Deep DuS-KFCM Segmentation with annotated images magenta area: Boundary by system (the result of proposed deep DuS-KFCM segmentation algorithm); green: Boundary by Gastroscopist (ground truth); White area: The fusion Image D with RA/RB as reference information

We start with taking each image from the training set and segment the gastric bleeding lesions from the image using DuS-KFCM fuzzy clustering technique. We repeat the above process for every image in the data set and use them to train our model – deep DuS-KFCM model. Then, for every testing image, we segment the GB lesion using the same method and then classify the lesion as benign or malignant using our model. The aim of this research is to get the contours of interest approximation in the medical images are determined by employing Dus-SKFCM and enhancement by deep learning model(region-growing).

The list of evaluation of performance and area for these images has been given in Table 3.

The one which evidently distinguishes the clusters is considered for segmentation. Experimental results show that the area segmented by the proposed method is clear and with high veracity. A direct visual comparison is shown in Figure 11.

We can observe that our method is robust for different GB image. the seed label image, which is generated automatically by deep learning algorithm, the magenta areas are foreground seed areas, and the black area are background seed area.

Experimentally, various kinds of performance analyses with state-of-the-art techniques are performed. Anyway, the proposed methodology outperforms better classification accuracy of 87.9475% than other methods. The results of proposed technique were compared with previously explored methods and are enlisted below, which is shown as Table 5.

**Table 5.**
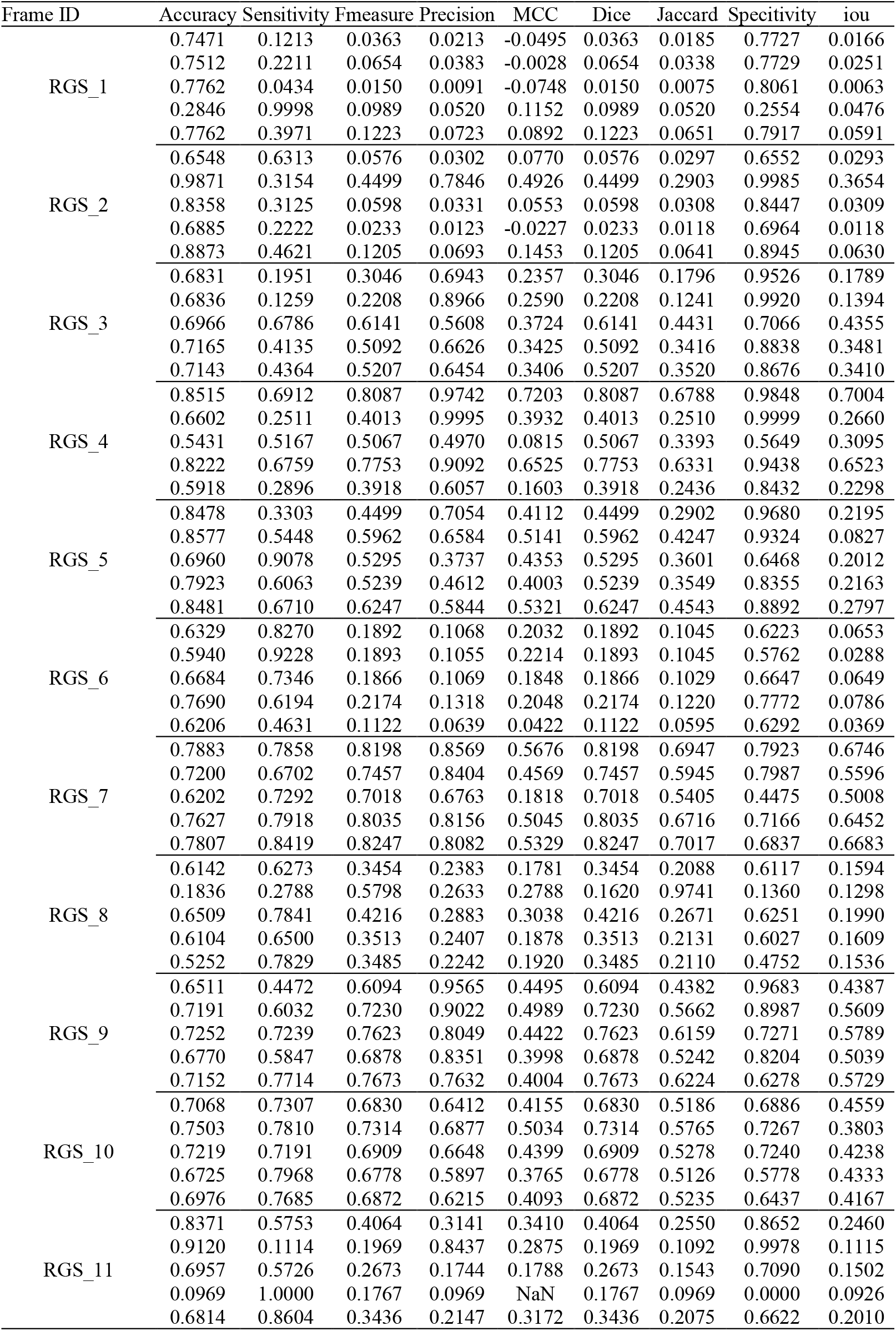
(Evaluation of Segmentation Method) Based on the performance on our challenging testing dataset.

## 7. Segmentation Accuracy

Accurate gastric bleeding lesion segmentation is an essential and crucial step for computer-aided gastric disease diagnosis and surgical planning. The performance evaluation of segmentation results shows that the proposed gastric bleeding segmentation method is accurate and efficient. Experimental results have been shown visually and achieve reasonable consistency. Accuracy obtained from the proposed work is 98% endoscopy gastric images.

The performance measures for segmentation of the deep DuS-KFCM method is analysed by accuracy of segmentation, specificity, precision and false measure (F-measure). dice-coefficient, and Jaccard-coefficient and ect. to evaluate quality of clustering.

Looking at Tables 5 for batch processing, it can be noticed that: the global accuracy, mean IoU, and mean F1 score results from the automatic segmentation’s performance evaluation, are all increased obviously after applying the preprocessing deep fuzzy enhancement step. This obvious quantitative enhancement assures the success of our proposed approach in batch automatic fuzzy segmentation. For making an overall view of the quantitative metrics over a GB images on deeplabv3+ (resnet50) based DuS-KFCM models, Tables 3 and 5 are merged and averaged producing a concentrated view for the quantitative batch frame work represented in Table 6. integrated into a novel fuzzy variational model for more accurate lesion delineation. The detection and segmentation performance evaluation performance metrics of state-of-the-art comparison based on segmentation results for each GB image are shown in Figures 12.

**Table 6.**
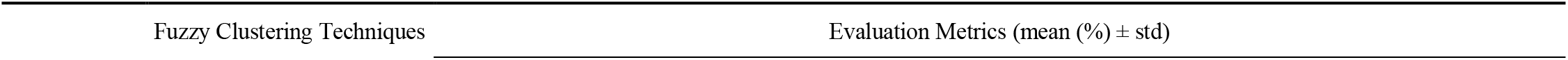
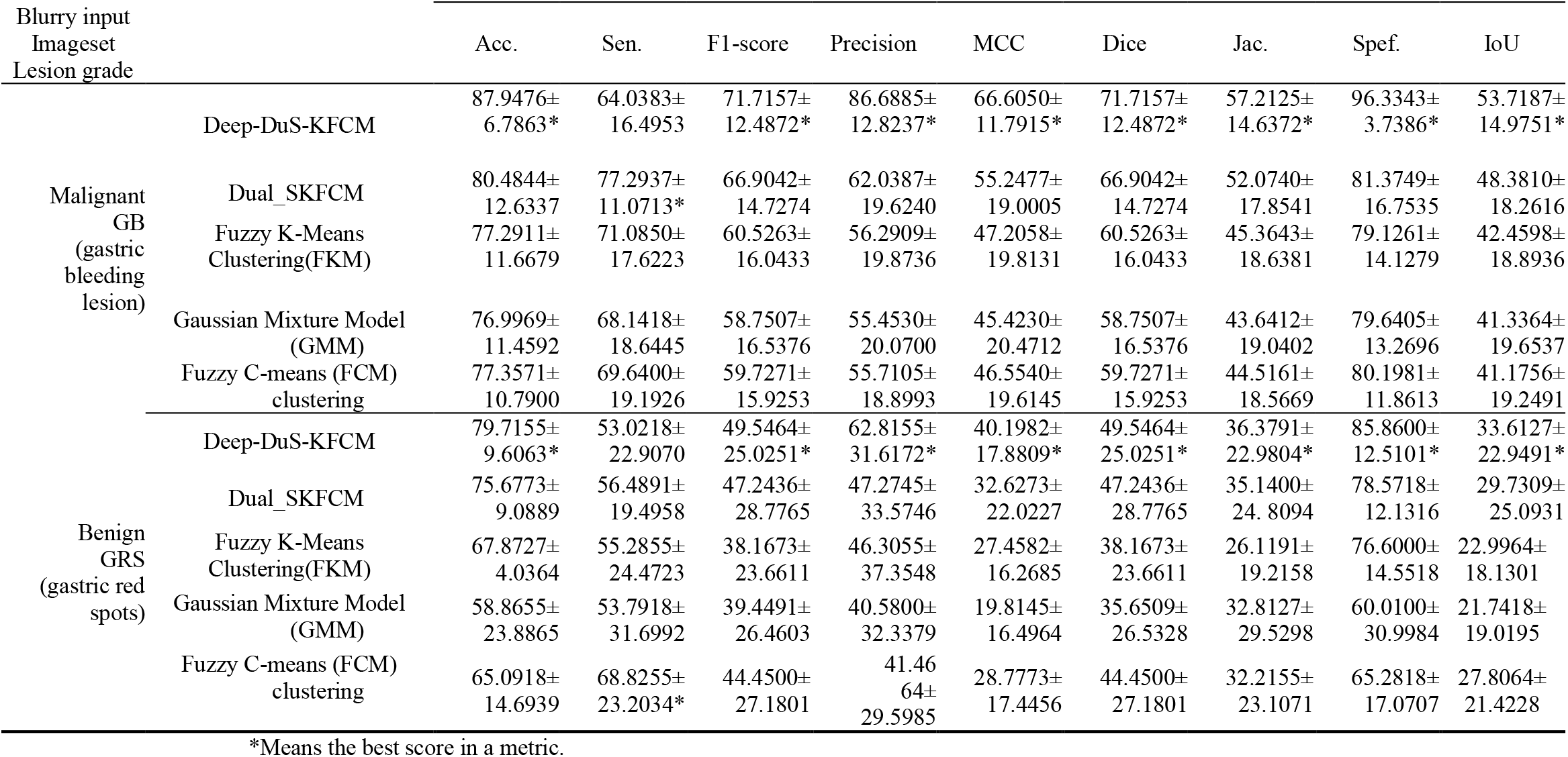
The Comparison of Clustering Algorithms DuS-FCM and Deep DuS-FCM for segmentation Gastric Blood Lesions using the mean and standard deviation of evaluation indexes.

**Figure 12.**
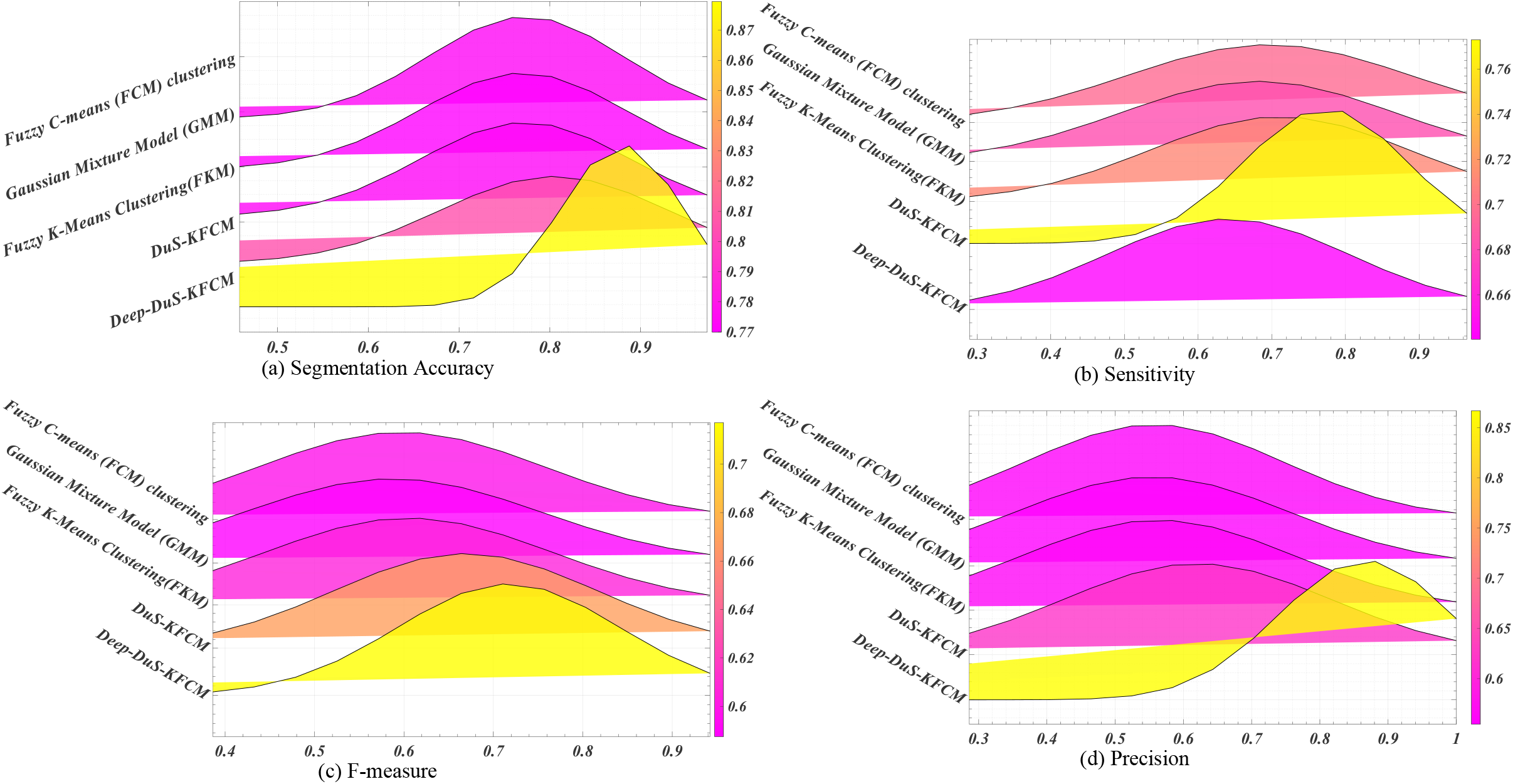
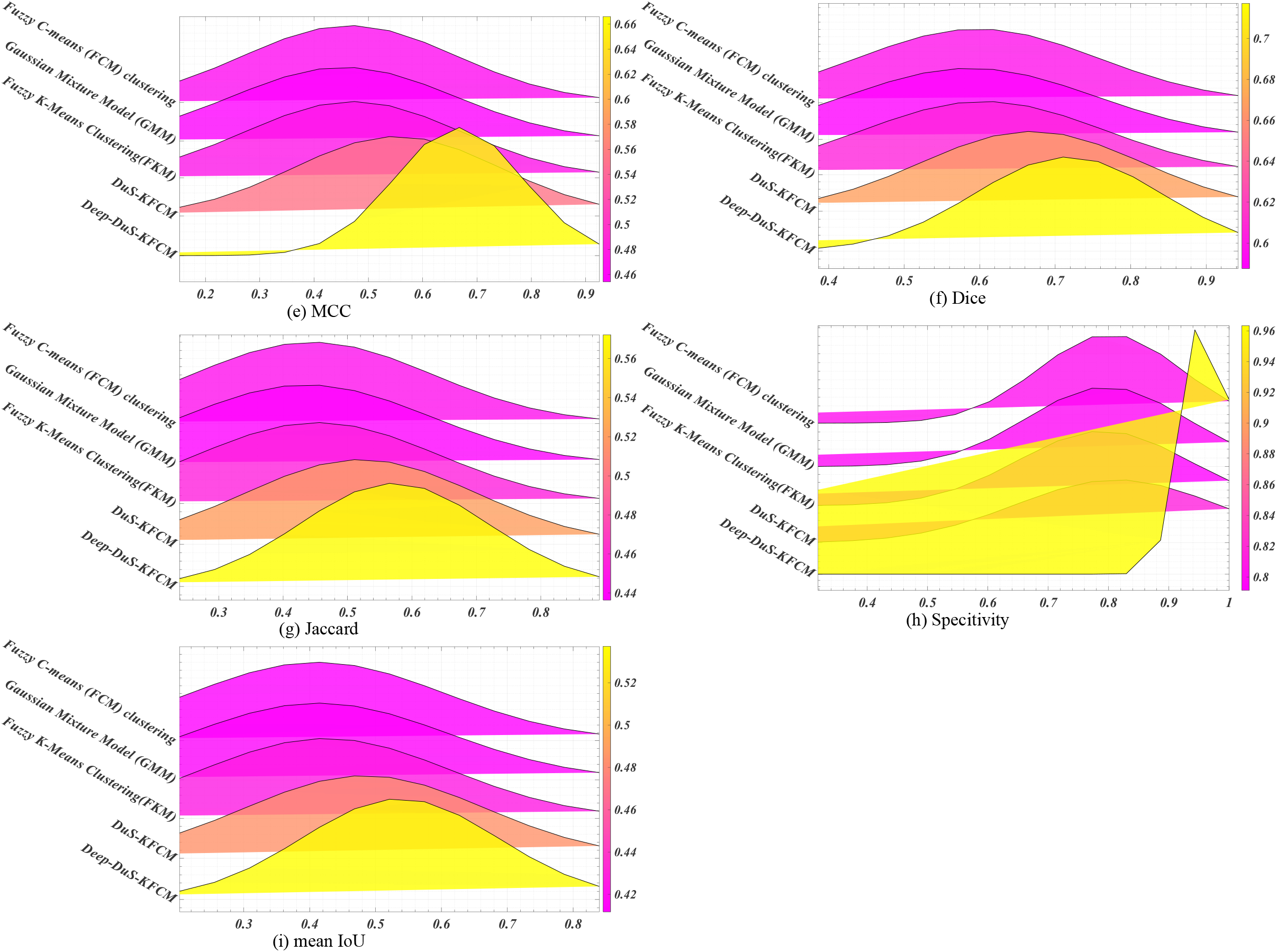
Comparing clustering models with 9 indexes via Gaussian Ridge Functions

Results are presented in Table 4. They indicate that the proposed method achieves significant improvements over other methods in terms of 9 evaluation metrics. Figure 1 and Figure 2 gives some visual comparison results of both the proposed method and other methods. Effect of predicted result in deep Du-SKFCM ensemble model reveal that the proposed method acquires a satisfactory balance between detailed information and contextual information. The evaluation results were better than those of existing methods.

We use criterion performance metrics including accuracy (ACC) sensitivity (Se), specificity (Sp), precision (prec.), matthews correlation coefficient (MCC), dice-coefficient, jaccard index, specificity(spec.) and intersection over union (IoU) to evaluate the classification performance. Description are defined as follows:

Accuracy determines that how many true positives TP, True negatives TN, False positive FP and False negatives FN were correctly classified. Segmentation Accuracy (SA) is defined as the ratio between the total number of classified pixels into the total number of pixels. To quantitative comparison of the segmentation performance, we use one objective indicator, i.e., the segmentation accuracy (SA) [7]. The specific score standard formula description of SA is computed as follows:

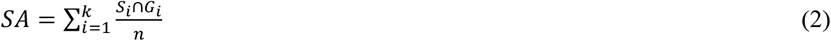

where the *S*_*i*_ represents a segmentation result, the *G*_*i*_ denotes the corresponding ground truth, the *k* is the number of clusters and *n* is the total number of pixels of images.

Sensitivity is the amount of positive items correctly identified:

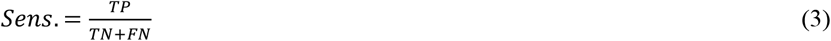

Precision: It checks how precise the model works by checking the correct true positives from the predicted ones. The precision is defined as the sum of TPs is divided by the sum of TPs and FPs. The performance of the precision is computed for segmented object, and it is capable of a returning only related instances to the classification model. The precision is expressed in Eq.4.

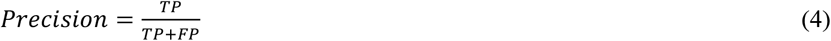

While, recall is defined as:

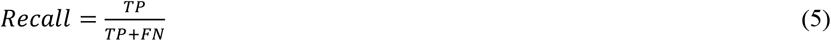

F1-score is the function of precision and recall. It is calculated when a balance between precision and recall is needed. F-measure is defined as the weighted harmonic mean of the precision and the recall is mathematically expressed in Eq.6.

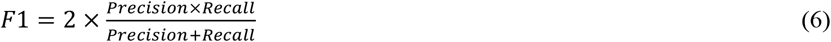

The Matthews correlation coefficient (MCC) is also used in our work. It is generally considered to be a more balanced indicator, even if the sample content of the two categories varies widely [8].

The Matthews correlation coefficient (MCC) is essentially a correlation coefficient between the actual classification and the predicted classification. Its value range is [-1, 1]. A value of 1 indicates a perfect prediction for the subject. A value of 0 indicates that the predicted result is not as good as the result of random prediction, and -1 means that the predicted classification is completely inconsistent with the actual classification. The MCC is defined as follows:

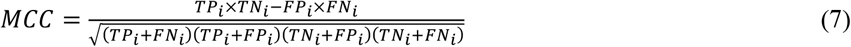

Dice-Coefficient: It is a statistical measure of similarity rate between two sample sets:

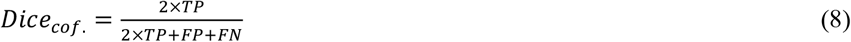

where TP is the number of true positive, FP is the number of false positives and FN is the number of false negatives.

Jaccard Index: It is a measure of similarity rate between two sample sets: Jaccard similarity: GT represents ground truth, and CS denotes cancer segmentation; Jaccard coefficient is defined as follows:

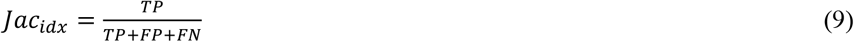

Specificity: It is the rate of correct identification of negative items: The precision is defined as the sum of TPs is divided by the sum of TPs and FPs. The performance of the precision is computed for segmented object, and it is capable of a returning only related instance to the classification models. The specificity is expressed in Eq 10.

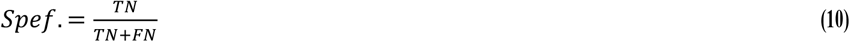

SEG score (object-wise Intersection over Union (IoU)) was used [9]. To calculate SEG, IoU (also known as Jaccard index) is needed for the calculation. IoU is defined as equation. The IoU is a very straightforward metric that is extremely effective. Simply put, the IoU is the area of overlap between the predicted segmentation and the ground truth divided by the area of union between the predicted segmentation and the ground truth. The intersection-over-union (IoU), also known as the Jaccard Index, is one of the most commonly used metrics in semantic segmentation. is one of the most commonly used metrics in semantic segmentation.

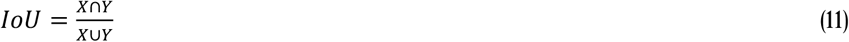

where X and Y are manually annotated segmentation mask and predicted segmentation mask, respectively. For every manually annotated object, the segmented object with the largest IoU is found.

Next, average IoU of all manually annotated objects is calculated. If IoU for any manually annotated object is smaller than 0.5, then IoU for this object is set to 0. This ensures that each manually annotated object can be paired with only one segmented object. The resulting SEG is an average of IoUs for all manually annotated objects.

## 8. Method verification and practical applications

In this section, to demonstrate the effectiveness of the proposed method deep DuS-KFCM, we compared with other recently reported prediction methods on the same gastric datasets. All the methods are performed using GRS cross-validation test. Tables details the comparison of the proposed method and other prediction methods on the gastric and red spot datasets, respectively. The quality of the clustering can be measured in two ways: directly or indirectly. In the direct method, we can apply some cluster validity measures to check whether the quality of the clustering is improving. In the indirect method, we cluster the data using the proposed method, and then the clusters are utilized to build the classifiers. The classifier performance is used as an indirect way to quantify the quality of the clustering. Note that this is possible only when the data are labeled. The system is vailed using gastric red-spots (GRS) images downloaded from various Web sites. We have retrieved RGS images from this dataset to provide a detailed comparison. The lesion of sample images are shown in Figure 13:

**Figure 13.**
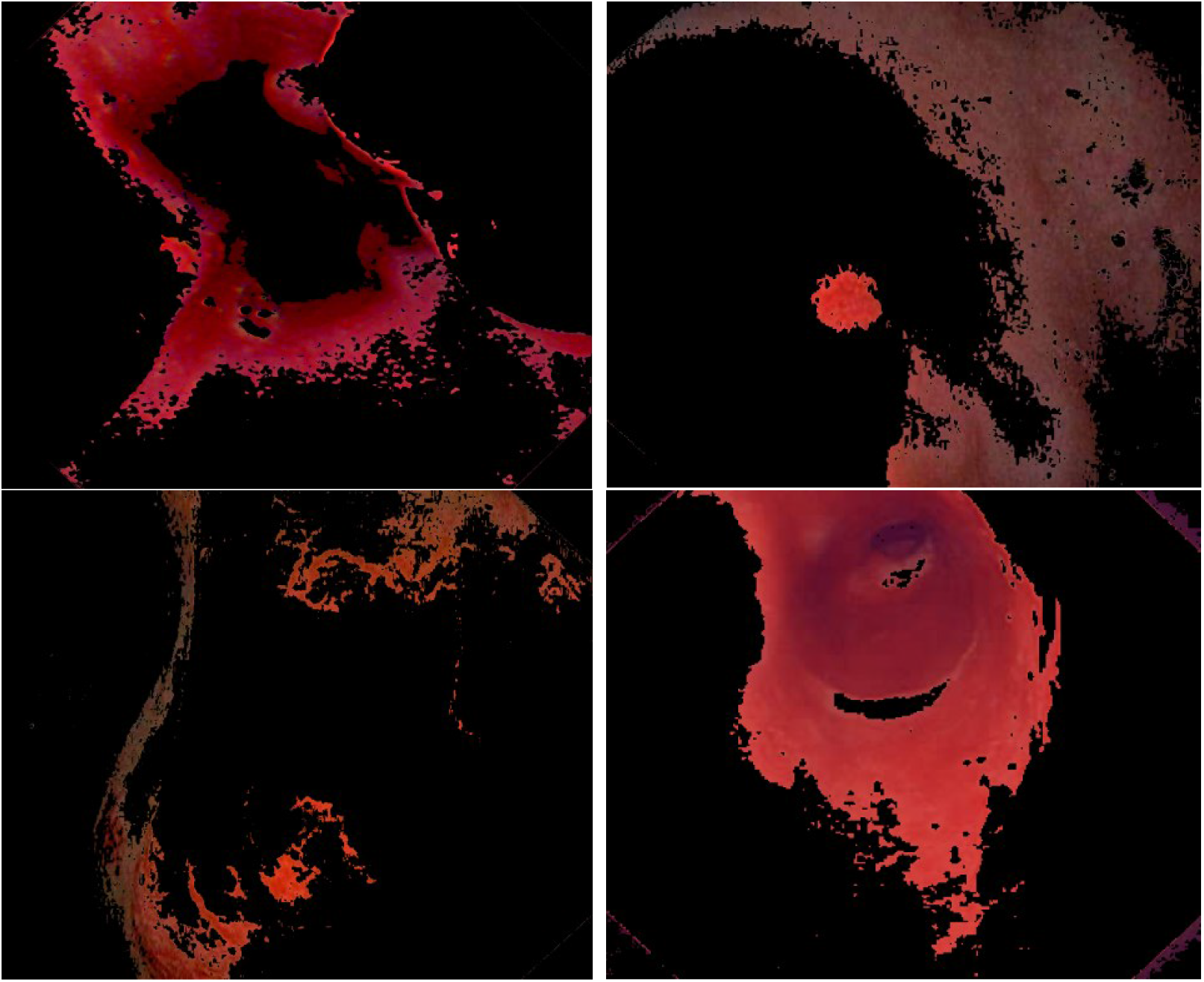
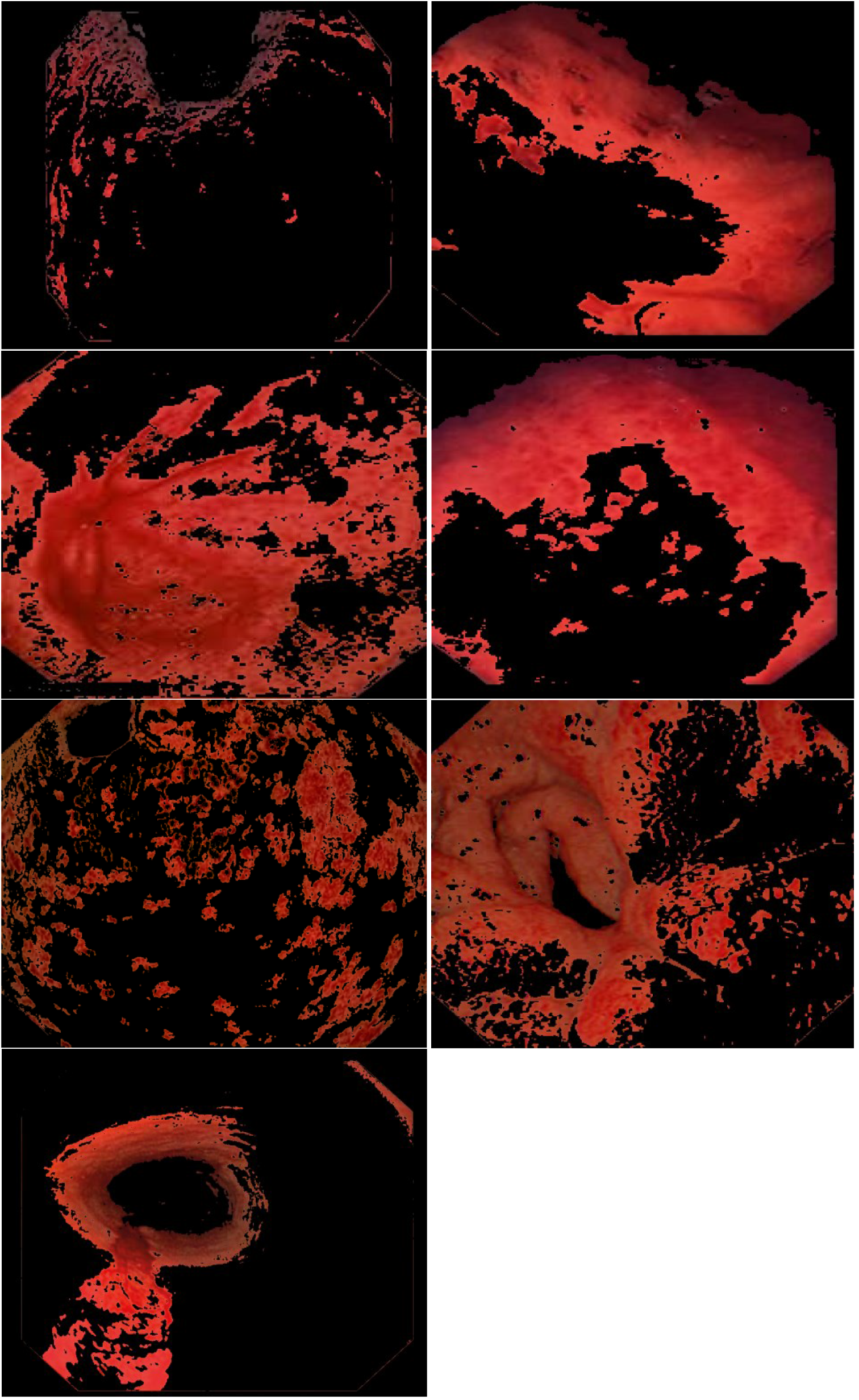
GRS Lesion Sample

The proposed work utilizes DuS-KFCM algorithm to extract the lesion from region of interest, where the deep DuS-KFCM input data are verified by using the GRS dataset in order to automate the process of segmentation. Visual comparison of the segmentation results on GRS images are shown in Figure. 14, which compare the segmentation results of the proposed fuzzy method with those obtained by GT on GRS dataset for each case. The magenta area is predicted by the proposed method; The green area is the manual area by Gastroscopist; The white area is true position area. This deep learning network was fed with only a small amount of GRS samples for learning in our study. Although the probability map could not accurately describe the lesions, it effectively distinguished the lesions from the complex background in endoscopy images.

**Figure 14.**
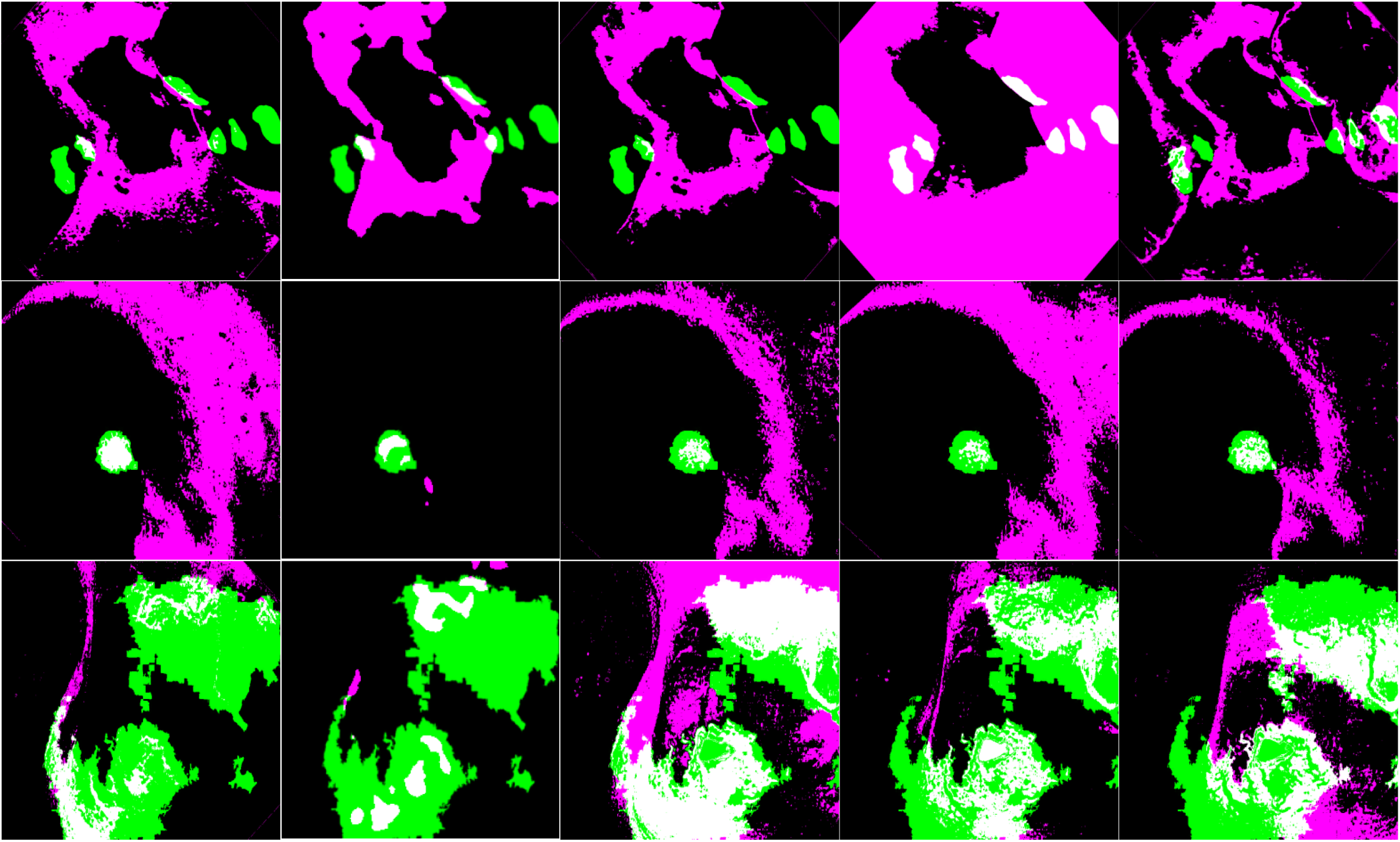
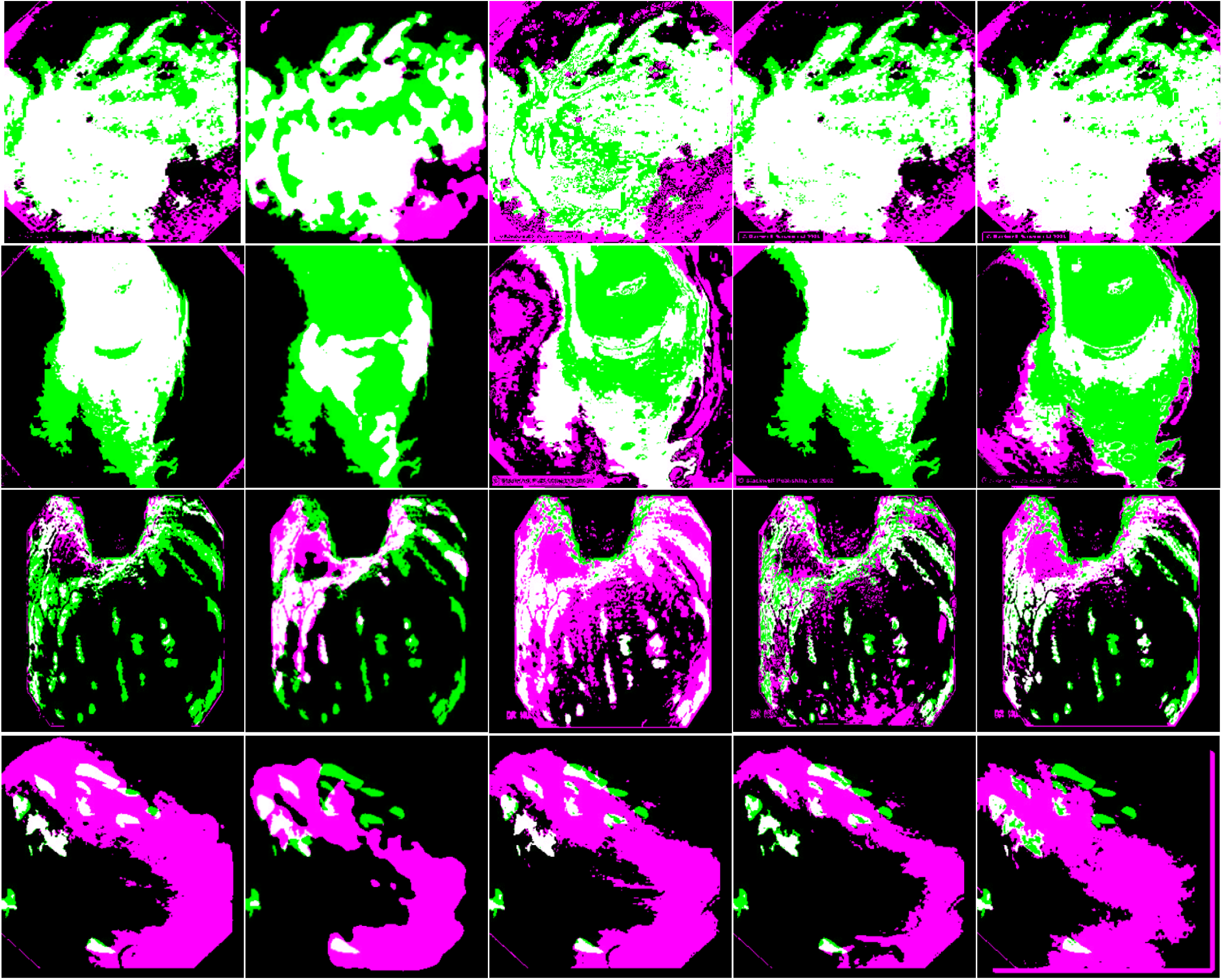
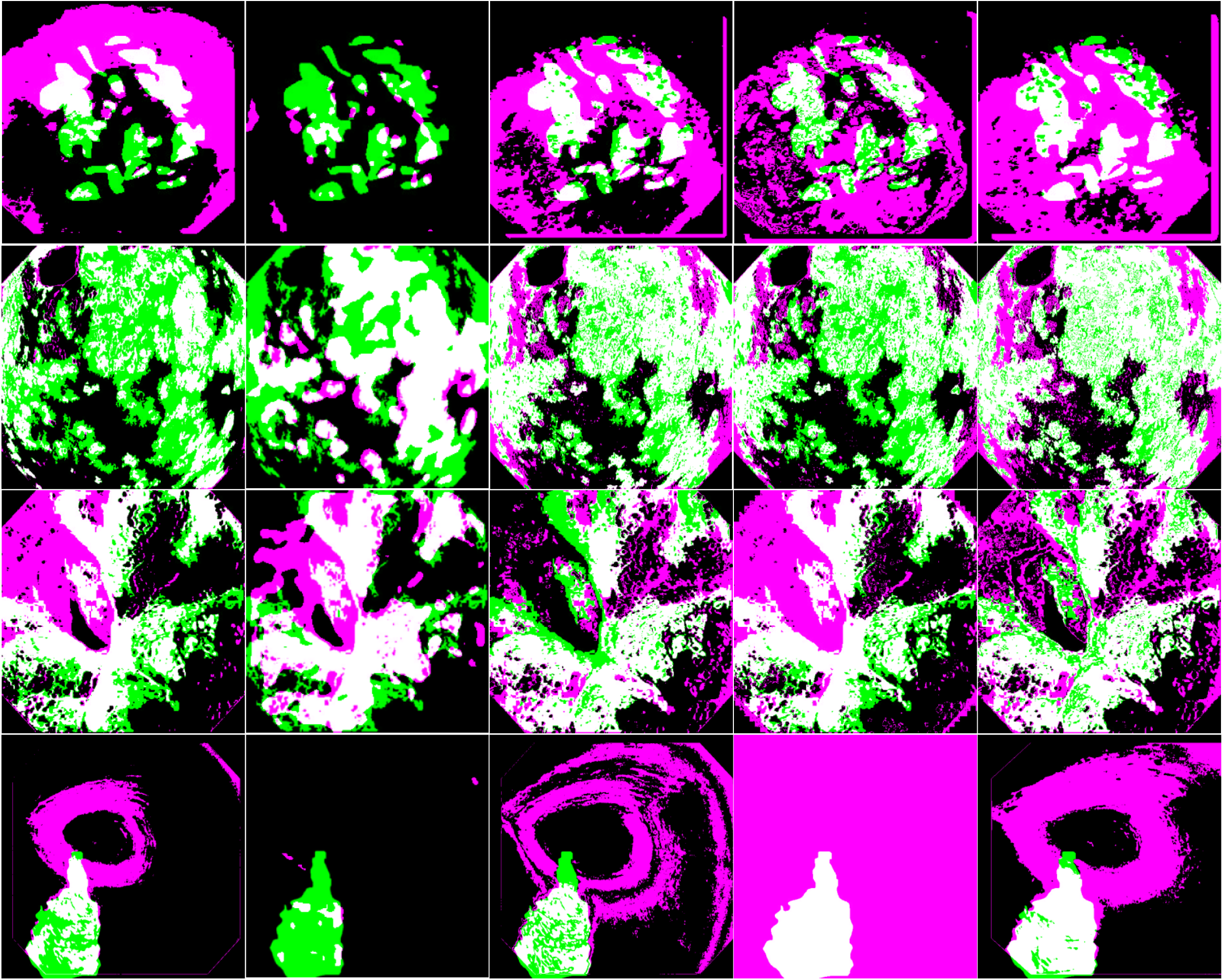
Results on the Pseudo-color Synthetic Images on GRS dataset: different fuzzy clustering (Proposed methods vs. Traditional threshold methods) leading to differently clustered images (Magenta area: Boundary by system; Green: Boundary by Gastroscopist(Ground Truth (GT)); White area: fusion Image D with RA/RB as reference information)

In terms of performance analysis of each object segmented image, to provide a detailed study, we have retrieved some more early gastric bleeding cancer images which is reported with the help of accuracy, precision, IoU and ect. in Table 5 and Figure. 14. For each input image, our method shows better robustness and identification to the lesion area which maintain the equal or nearly equal high recognition rate for each.

As Figure 15, we compared the image segmentation quality scores with some related works. This experiment validates the segmentation performance of the proposed algorithm with other state-of-the-art methods including the DuS-KFCM (we proposed), Fuzzy K-Means clustering (FKM), Gaussian Mixture model (GMM) and Fuzzy c-means clustering (FCM) method. While compared to the existing methods, the proposed novel fuzzy set segmentation algorithm demonstrates better segmentation results. Experiments and Results Discussions

**Figure 15.**
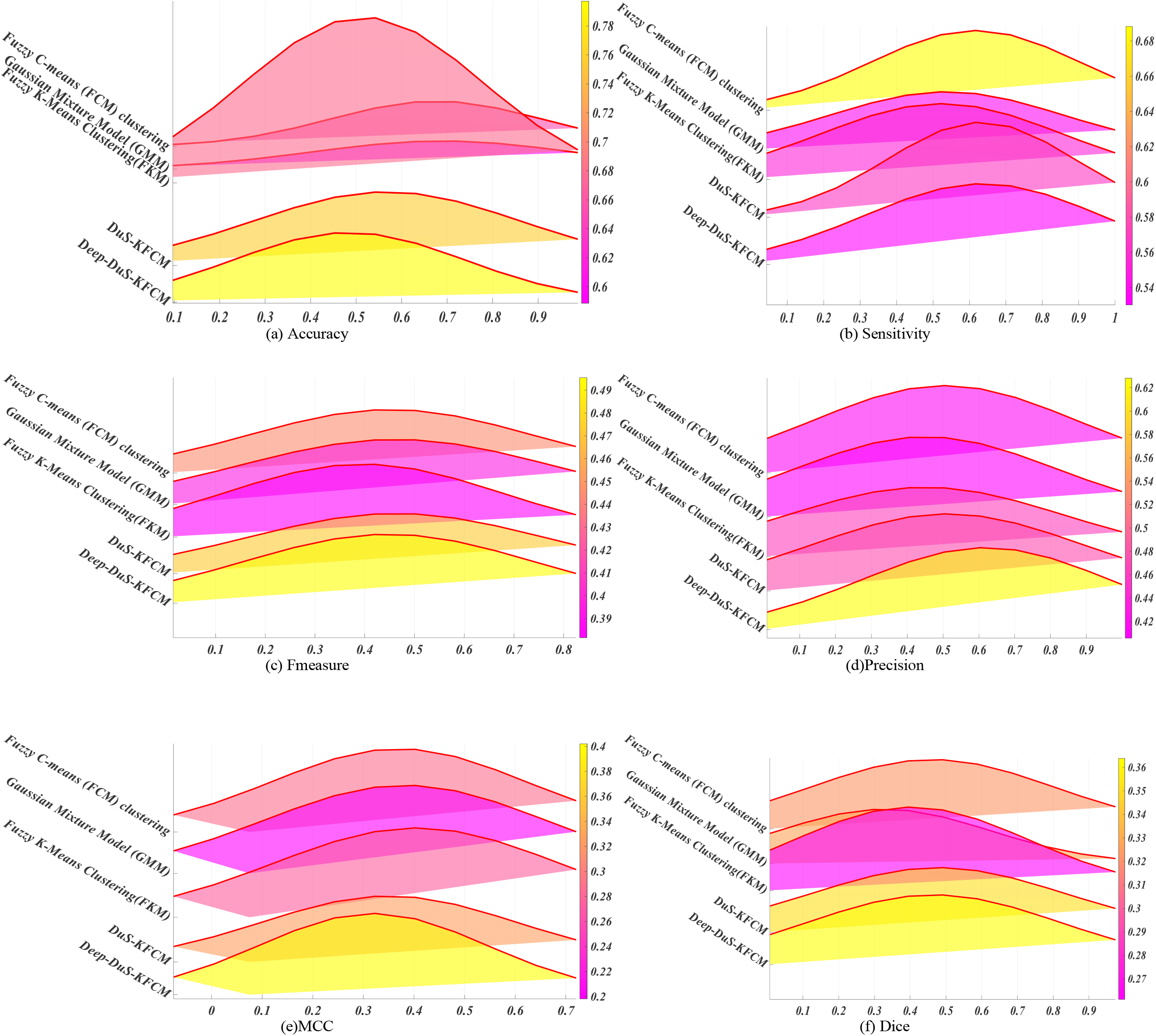
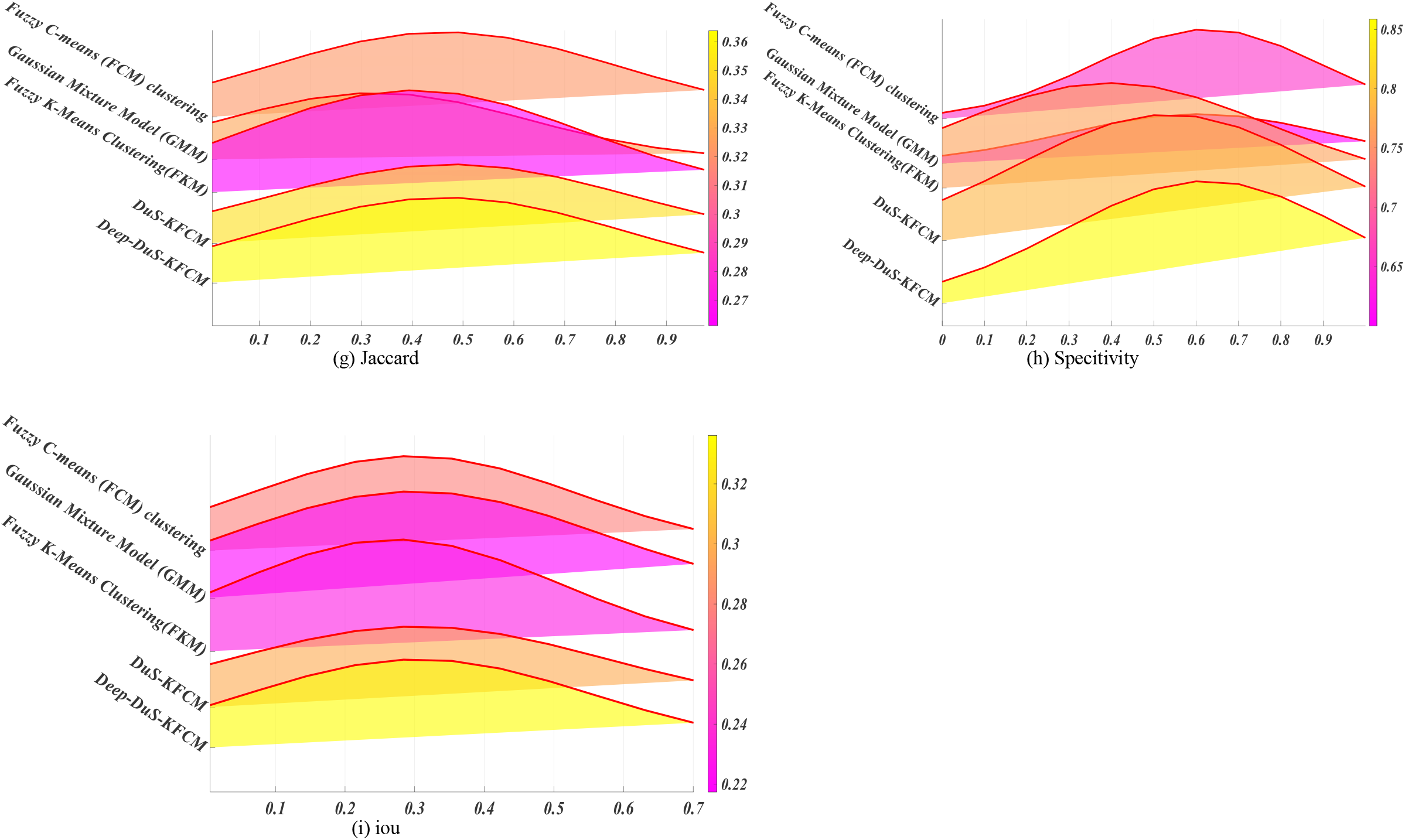
Assessment of the reliability of clustering models on GRS dataset by different methods

In this subsection, we show the effectiveness of our method on some simple segmentation tasks. These images have obvious different colors and have a reasonable criterion of the fuzzy segmentation, Average performance results for several different strategies to segment is shown as the Table 6.

The performance of the proposed algorithm is evaluated on medical images from different modalities. The results confirmed its effectiveness for medical image segmentation. With different quantization schemes for all databases, experimental results indicate that deep DuS-KFCM provide best accuracy, which has better recognition accuracies as compared to other method. Performance of the model tested using the former dataset achieved Avgacc. 53.7157; whilst, using the later dataset performed Avgacc 33.6127, which are the highest performance from the other tested models. The deep DuS-KFCM are observed to improve the recognition average precision of 2nd experiment from around 62.8155 to 86.6885.

The proposed deep DuS-KFCM is evaluated in terms of Recall, Specificity, Precision and F-measure and the experimental results showed that the proposed method improved the system performance from 0.3 % to 14.35 % compared to existing methods.

After incorporating them together, the mIoU of the generated pseudo labels was further promoted to 53.7187% on the train set and 33.6127% on the val. set. The applied work on small dataset the output evaluation metrics of the batch mode for all SS operation on both benign and malignant GB images by five fuzzy segmentation models.

Average Accuracy: It is a measure of effectiveness of the classifier: where m is the total number outputs of the system.: The average accuracies (OA%) of each technique

Where X is the total number of medical images.

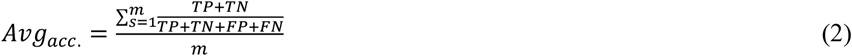

AP (Average precision) is a popular metric in measuring the accuracy of object detectors. The average precision rate may be calculated as follows:

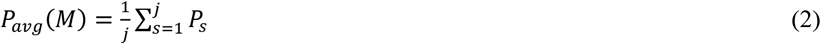

In eq. 18 Pavg(M) represents the average precision rate for category (M), where j is the total no of images in that category.

Mean intersection over the union (MIoU): MIoU refers to the mean IoU of the image calculated by taking the IoU of each class and averaging them. The MIoU is positively correlated with the segmentation effect of the model, and because of its simplicity and representativeness, it is often used as the main figure of merit to evaluate performance of image semantic segmentation models.

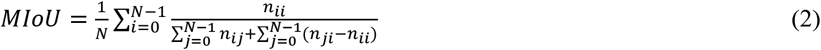

where is the number of pixels whose class semantics are recognized as class, namely, FN (false negatives). The MIoU is positively correlated with the segmentation effect of the model, and because of its simplicity and representativeness, it is often used as the main figure of merit to evaluate performance of image semantic segmentation models.

## 9. Conclusions and Perspectives

In this paper, we developed a novel method for automated detect the localization of masses in gastric bleeding images. a new coarse-to-fine method is proposed to segment gastric bleeding lesion for abdominal Esophagogastroduodenoscopy (EGD) images, called deep DuS-KFCM. The comparisons between the results produced by the proposed method and the manually segmented images provided by GB database showed that the proposed method can accurately and automatically extract the masses from ROI, which is comparable to an expert analysis and superb to available cancer segmentation methods.

The proposed deep DuS-KFCM method can be beneficial to developing a system for clinical images used for diagnoses. In this method, three types of images such as endoscopy results for gastric bleeding cancer, normal, and gastric red-spots image. The proposed method achieved 87.9476% and 79.7155% for the accuracy coefficient whereas 86.6885% and 62.8155% for the precision coefficient, respectively. The accuracy and precision coefficients are utilized to compare the diversity and similarity pixels between the images. The proposed method indicates a favorable accuracy for cancer detection through each medical image segmentation that we used in a computer-aided diagnosis. The computational costs are disregarded because the detection rate has reached a favorable degree. The accuracy degree for detecting cancer in patients through the medical image segmentation is promising. This method can be utilized to classify the type of cancers in accordance with the clinical diagnosis method.

In the future, there is still space for improvement in the image analysis of gastric cancer. Future work is planned to optimize our algorithm to be able to detect the non-brain original cancer metastasis. In addition to medical problems, we intend to apply the proposed technique to cluster the microarray genomic data, where missing values are encountered quite often due to the limitations of the experiments. there is a lack of complete, clear, and accurately labeled pathological images, so the establishment of more perfect data can provide great help for the experiment. Finally, in terms of feature extraction and classifier design, there are many novel techniques that can be applied to gastric cancer image analysis, which is a promising and valuable research direction.

The bleeding area error was mainly caused by occlusion, an inherent limitation with 2D images. Therefore, it will be necessary to extract traits from 3D images in the future.

Since the full set of features extracted and selected in this manuscript has reached a very good characterization of gastric bleeding areas, it could be interesting to apply them over other types of abnormal gastric blood lesion---(GRS)gastric red spots to detect bleeding morphologic changes of some gastric diseases.

It would be interesting to apply the methodology developed to assess their effectiveness in the classification of abnormal gastric blood areas in different types of cases.

Since the DuS-KFCM clustering algorithm can work with color spaces, the potential applications are not only limited to gastric bleeding images obtained from gastric endoscopy dataset but to other medical application areas, which require segment images with different regions with color threshold and spatial similarities.

Although the DuS-KFCM segmentation experiments were implemented with two clusters (normal & GB), the algorithm can be used with more clusters, increasing the details in the regions or the number or regions obtained.

The rich capillary zone around the blood has been explored only to calculate an approximated measure of the capillary blood projections. It would be interesting to extract more features from this region to analyze their contribution to the characterization of the gastric bleeding area.

## Notes

### Competing Interest Statement

The authors have declared no competing interest.

